# Energy landscape steering mediates dynamic coupling in ATP-driven protein translocation by the bacterial Sec machinery

**DOI:** 10.1101/793943

**Authors:** Joel A. Crossley, William J. Allen, Daniel W. Watkins, Tara Sabir, Sheena E. Radford, Roman Tuma, Ian Collinson, Tomas Fessl

**Author notes:** CytoSeek, Science Creates Old Market, Midland Road, Bristol, BS2 0JZ (Daniel W. Watkins).

## Abstract

The Sec translocon is a highly conserved membrane complex for transport of polypeptides across, or into, lipid bilayers. In bacteria, the core protein-channel SecYEG resides in the inner-membrane, through which secretion is powered by the cytosolic ATPase SecA. Here, we use single-molecule fluorescence to interrogate the dynamic state of SecYEG throughout the hydrolytic cycle of SecA. We show that the SecYEG channel fluctuates between open and closed states faster (*∼*20-fold during transport) than ATP turnover; while the nucleotide status of SecA modulates the rates of opening and closure. Interestingly, a SecY variant (PrlA4), exhibiting faster protein transport, but unaffected ATPase rates, increases the dwell time in the open state, facilitating pre-protein diffusion through the pore; thereby improving the efficiency of translocation. Thus, contrary to prevailing structure-based models, SecYEG plays an integral part in the translocation mechanism through dynamic allosteric coupling in which SecA ‘steers’ the energy landscape of the protein-channel.

## Introduction

A fascinating class of biological molecular machines are those operating upon biopolymer substrates, converting chemical energy derived from ATP binding and hydrolysis into cycles of conformational changes and mechanical work. Examples of these molecular motors include the helicases, unfoldases, chromatin remodelling complexes, primary membrane transporters, protein degradation assemblies^1–3^, and the subject of this paper–the protein translocases, exemplified by the ubiquitous secretory (Sec) machinery.

The bacterial translocon is minimally composed of the integral inner-membrane core-complex SecYEG, and a peripherally associated cytosolic motor ATPase, SecA (Fig. 1 and Supplementary Fig. 1). The complex of the two (SecYEG:A) is necessary and sufficient for the translocation of unfolded polypeptides across the inner membrane^4, 5^. Transport substrates of the Sec machinery–periplasmic and outer membrane proteins (OMPs)–are transported post-translationally through SecYEG:A as pre-proteins with N-terminal cleavable signal sequences (SS)^5, 6^. SecYEG also directly associates with ribosomes and mediates co-translational insertion of inner membrane proteins^7^. The mechanism by which SecYEG adapts and performs these various tasks remains unresolved^8–11^. This core-reaction is facilitated by ancillary factors to improve the efficiency of secretion and insertion^12–16^.

**Fig. 1.**
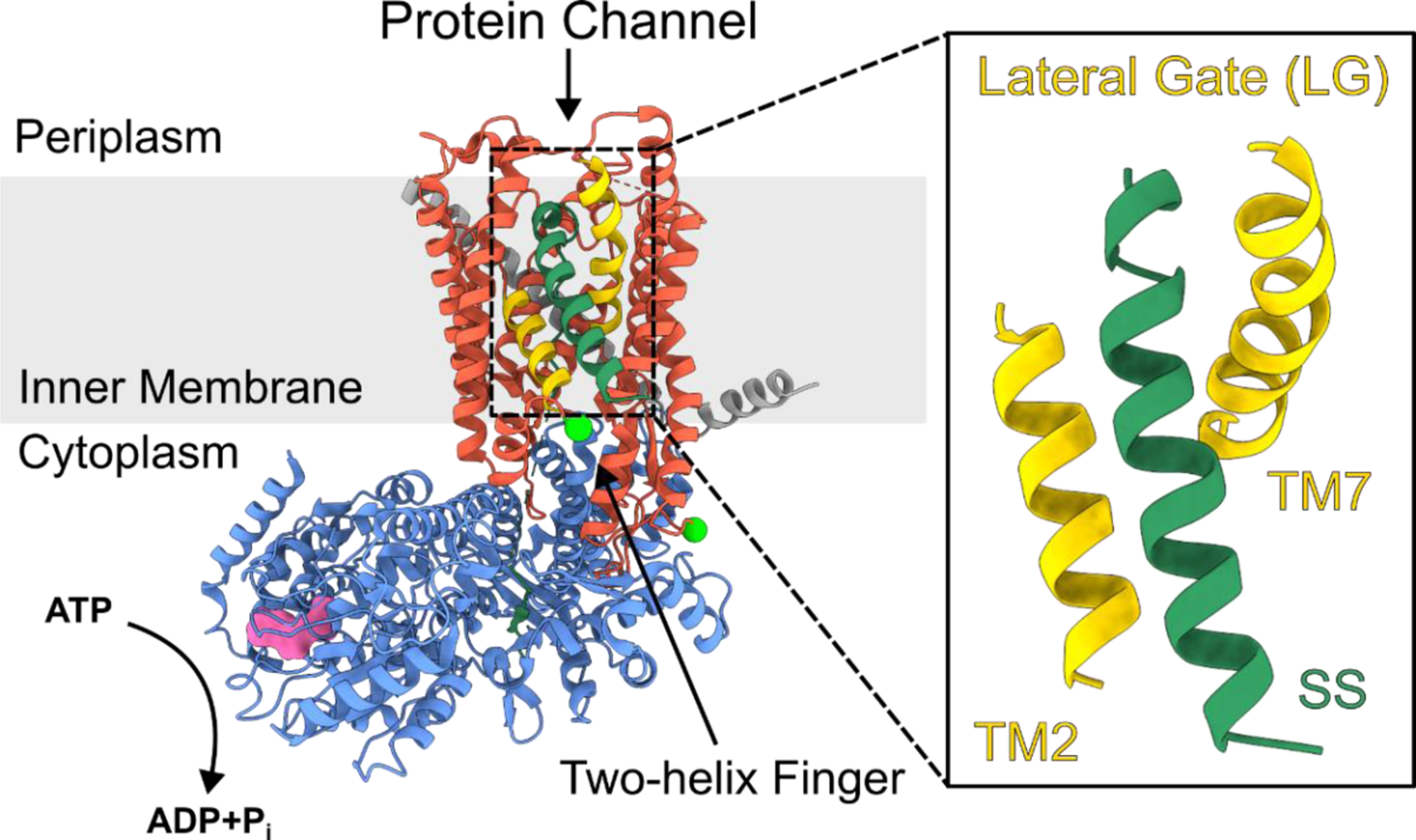
The electron cryo-microscopy structure of the SecYE:A translocon (PDB: 6ITC^17^, structure determined in the absence of SecG). SecY is shown in red, SecE in dark grey and SecA in blue. The two transmembrane helices that comprise the lateral gate (LG) are shown in yellow (TM2 + TM7). The inner membrane is shown in light grey. The signal sequence (SS) is shown in dark green. ADP is shown in surface representation (pink) within the nucleotide binding pocket (more details of SecA structure are shown in Supplementary Fig. 1). The positions of the mutations used to attach the fluorescent labels used in this work are shown as bright green spheres (A103C and V353C). Note that dye attachment at each site is stochastic with the labelling protocol used in this study.

The protein channel is formed through the centre of SecY^17^, adjacent to a lateral gate (LG) between transmembrane helices TM2 and TM7, which opens to the bilayer^18^ (Fig. 1); opening of which is required for pre-protein translocation^19, 20^. Protein transport is driven by the associated SecA, a DEAD-box ATPase with two nucleotide binding domains, which together form a single ATP binding site^21^. The two-helix finger (2HF) domain of SecA has been proposed to act as a sensor regulating nucleotide exchange^8, 22^, or alternatively, to directly push the translocating polypeptide across the membrane^9, 10, 23^.

The precise nature of the protein translocation mechanism through SecYEG:A has divided opinion, owing to the inherent complexity of the system^8–11^. As for other molecular machines which convert the chemical potential of nucleoside triphosphate hydrolysis into directional motion, there are two limiting cases of energy transduction: power-stroke and Brownian ratchet^24^. The former involves deterministic or direct coupling, in which each conformation imposed by the stage of the nucleotide hydrolysis cycle is linked with a well-defined conformation of the effector part (*e.g.* a mechanical lever or substrate binding site). On the other hand, a Brownian ratchet mechanism exhibits loose coupling between the conformations of the nucleotide-binding site of the motor ATPase and the effector part^25^. Both cases can be illustrated in terms of a simplified energy profile along the mechanical reaction coordinate (Supplementary Fig. 2). The power-stroke profile features deep minima at either the pre-stroke or post-stroke position which directly correspond to the nucleotide state of the ATPase. In the Brownian ratchet case, the nucleotide hydrolysis cycle biases a shallow energy profile (*i.e.* with low energy barriers between states) towards certain conformations, which in turn undergo rapid interconversion between the available states due to thermal fluctuations.

Power-stroke mechanisms have been demonstrated for many cytoskeletal motors^26^, while Brownian ratchet schemes have been implicated for other ATP driven systems, such as the ClpX polypeptide unfoldase^27^. Both mechanisms have been considered for the SecYEG:A translocon: the power stroke model invokes a large, piston-like motion of the 2HF in SecA, imparting force directly to the substrate, with SecYEG considered a passive pore^9, 10, 23^. Alternatively, SecYEG has been proposed to support a Brownian ratchet, allosterically communicating with SecA^8^. A recent single-molecule Förster resonance energy transfer (smFRET) study detected conformational changes of the 2HF taking place on timescales (measured as dwell times in different states) of 100 - 400 ms^10^. This is similar to the timescale of the ATP hydrolysis cycle^28, 29^ (∼100 ms) and was interpreted as evidence that SecA acts *via* a directly coupled power stroke. However, in all published structures of SecYEG:A complexes obtained to date^17, 30–33^ there is less than 1 nm available for 2HF movement, not enough for the proposed power stroke. Moreover, covalent crosslinking of the 2HF to SecYEG does not prevent translocation activity^22^, suggesting that the 2HF does not need to move long distances during protein transport.

A recent structural study of SecYE (structures determined in the absence of SecG) and SecA found no movement of the 2HF between two nucleotide occupancy states (ADP·BeF_3_^−^ and ADP)^33^. Based on these static structures, the authors proposed that two SecA loops in the pre-protein cross-linking domain (PPXD) and the nucleotide binding domains (Supplementary Fig. 1) respond to ATP binding and hydrolysis in a fashion similar to monomeric RecA-like helicases and effect directional motion towards the SecYEG channel, once again considered as a passive pore^33^. However, evidence suggests that SecYEG is far from a static bystander. Indeed, it is well known that the SecYEG channel is actively gated by SecA: biochemical data and molecular dynamics simulations show that the opening and closure of the LG is linked to the nucleotide state of the associated SecA^8, 34^. Consistent with this, nucleotide turnover of SecA is affected by SecYEG: i) the SecA ATP hydrolysis rate increases ∼27-fold when associated with SecYEG, and ∼760-fold during translocation^28^; and ii) ADP release is affected by the LG conformation^8^. Further evidence for two-way allosteric coupling between the channel and motor components was provided by hydrogen-deuterium exchange experiments^34^.

A better understanding of the coupling between SecA and SecYEG is key to the reconciliation of these contrasting mechanisms. Given the dynamic nature of SecYEG^35^, high resolution static structures are not able to distinguish between them: the dynamic motions in SecA and SecYEG are undiscernible, and fleetingly populated states may be missed. To address this question, we employ smFRET analysis of the dynamic motions of the SecYEG channel throughout the SecA ATPase cycle. Our study also provide quantitative analysis of dwell times in the different states observed using multi-parameter photon-by-photon hidden Markov modelling (mpH^2^MM)^36^. We show that SecYEG undergoes transitions between open and closed states on a sub-millisecond time scale, much faster than SecA-catalysed rate of ATP hydrolysis (> 100 ms)^10, 28^. Nonetheless, the rates of these transitions are controlled by the nucleotide occupancy of SecA. Hence, rather than a direct coupling of SecA ATP binding/ hydrolysis to SecYEG channel motions, the SecA ATPase cycle regulates SecYEG opening and closing by modulating its underlying shallow energy landscape. Quantitative analysis of the state dynamics also revealed a previously undetected, post-hydrolysis, committed state– linked to protein translocation, but preceding the equilibrium ADP-bound state. Finally, we show that rapid translocation of the poorly selective SecY variant PrlA4 (Supplementary Fig. 3) correlates with perturbed channel motion, consistent with SecYEG dynamics being a key factor in the process of polypeptide transport.

## Results

### Intrinsic dynamics of SecYEG observed through interconversion between open and closed conformations

To monitor the conformation of the SecYEG we employed a previously engineered version of SecY (SecY_A103C-V353C_EG) which contains two unique cysteines introduced into TM2 and TM7 (Fig. 1). This pair, when labelled with fluorescent dyes, was demonstrated as a sensitive reporter of opening and closure of the channel and LG^8^. Previously, we identified at least two states, open and closed, based on smFRET measurements. Another FRET signal was observed arising either from a genuine third state (partially open), or from an average of fast interconversion of the open and closed states. At the time, these possibilities could not be resolved with the camera-based single molecule setup (200 ms per frame)^8^.

Here, we deployed a single-molecule diffusion-based confocal setup, allowing for the detection of conformational dynamics on the sub-millisecond timescale^37^. The implementation of pulsed interleaved excitation^38^ provides access to E_raw_ (raw FRET efficiency) and S_raw_ (raw stoichiometry) enabling the use of mpH^2^MM^36^ for detection of transitions and states, and for extraction of the corresponding dwell times (Fig. 2c; Supplementary Methods and Supplementary Table 1). mpH^2^MM also effectively deals with contributions from molecules with incomplete fluorophore labelling or those undergoing undesirable photophysics which can interfere with analysis of dynamic populations (*e.g.* donor/ acceptor-only species and dark donor/ acceptor species arising as a result of fluorophore ‘blinking’).

**Fig. 2.**
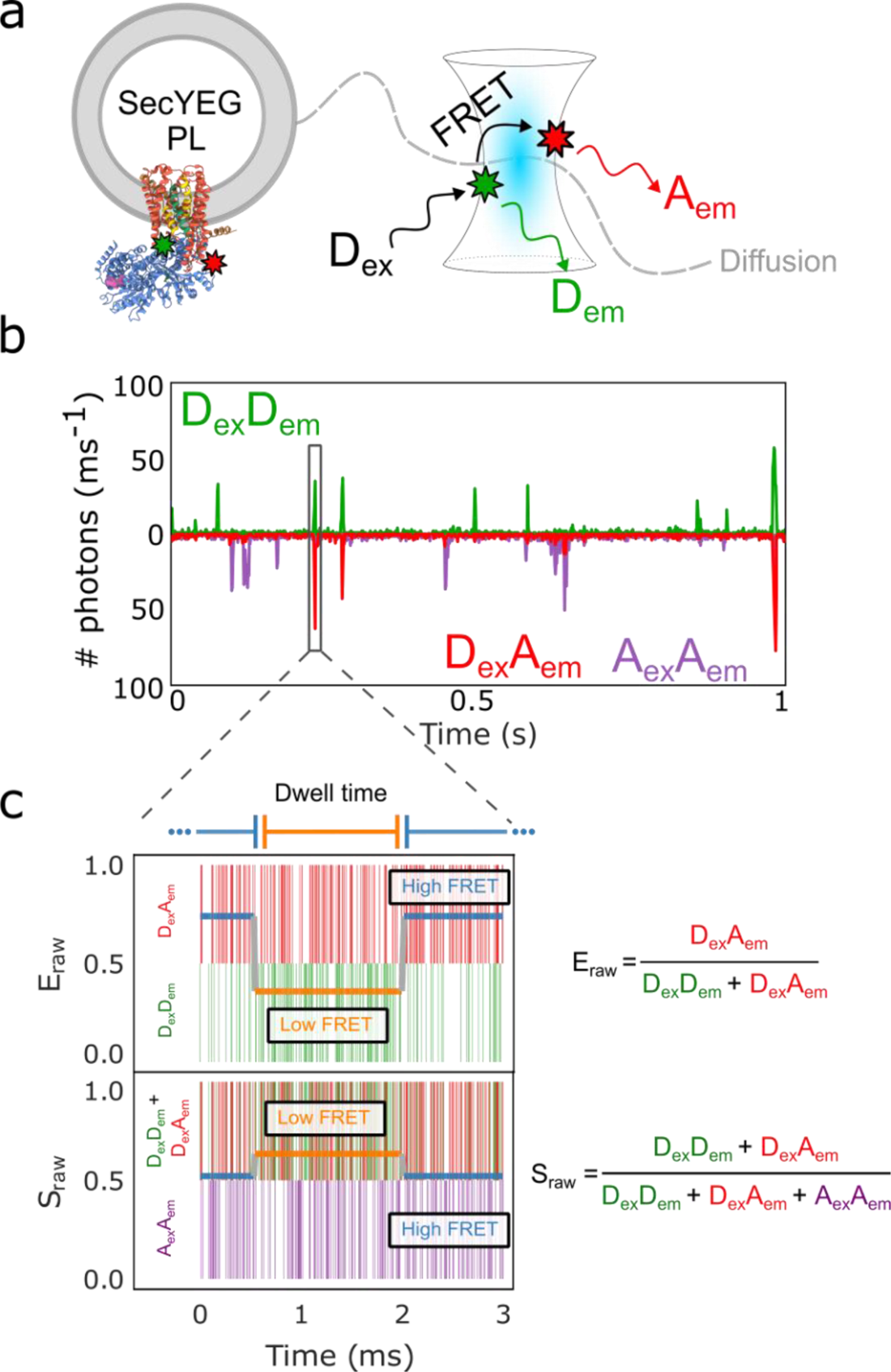
Single-molecule FRET methodology. a) Confocal volume with illustration of a single proteoliposome (PL) with embedded SecYEG diffusing in and out of the confocal volume alongside. The positions of the dyes are indicted using green/red stars. Note that SecYEG is shown in a single orientation here, but inserts equally in both orientations into the liposomes under the conditions used^63^. Hence only ∼50% of molecules will bind SecA in the experiments presented (see Methods). b) A representative single-molecule time trace showing photons from the D_ex_D_em_ (Donor Excitation, Donor Emission) in green, D_ex_A_em_ (Donor Excitation, Acceptor Emission) in red, and A_ex_A_em_ (Acceptor Excitation, Acceptor Emission) in purple. c) A single-molecule photon time trace of a single burst. Photons are represented by vertical bars. The most likely state path as identified by the Viterbi algorithm as derived by mpH^2^MM is overlaid as two horizontal coloured lines in both E_raw_ and S_raw_.

The approach for dual labelled SecYEG alone (i.e. in the absence of SecA) in liposomes reconstituted from *E. coli* polar lipid is described in Fig. 3. mpH^2^MM followed by the modified Bayes Information Criterion (BIC’) analysis (see Supplementary Methods) indicated that the datasets are best explained by four classes (Fig. 3a). Two of them being FRET states associated with conformations of SecYEG designated as open and closed. This is based on respective low and high FRET efficiency, consistent with the known distances between the two labelled sites in representative crystal structures of the two states (Fig. 3b,c). The remaining two classes represent contributions from the donor only (or dark acceptor) and acceptor only (or dark donor) species, and are therefore not physiologically relevant.

**Fig. 3.**
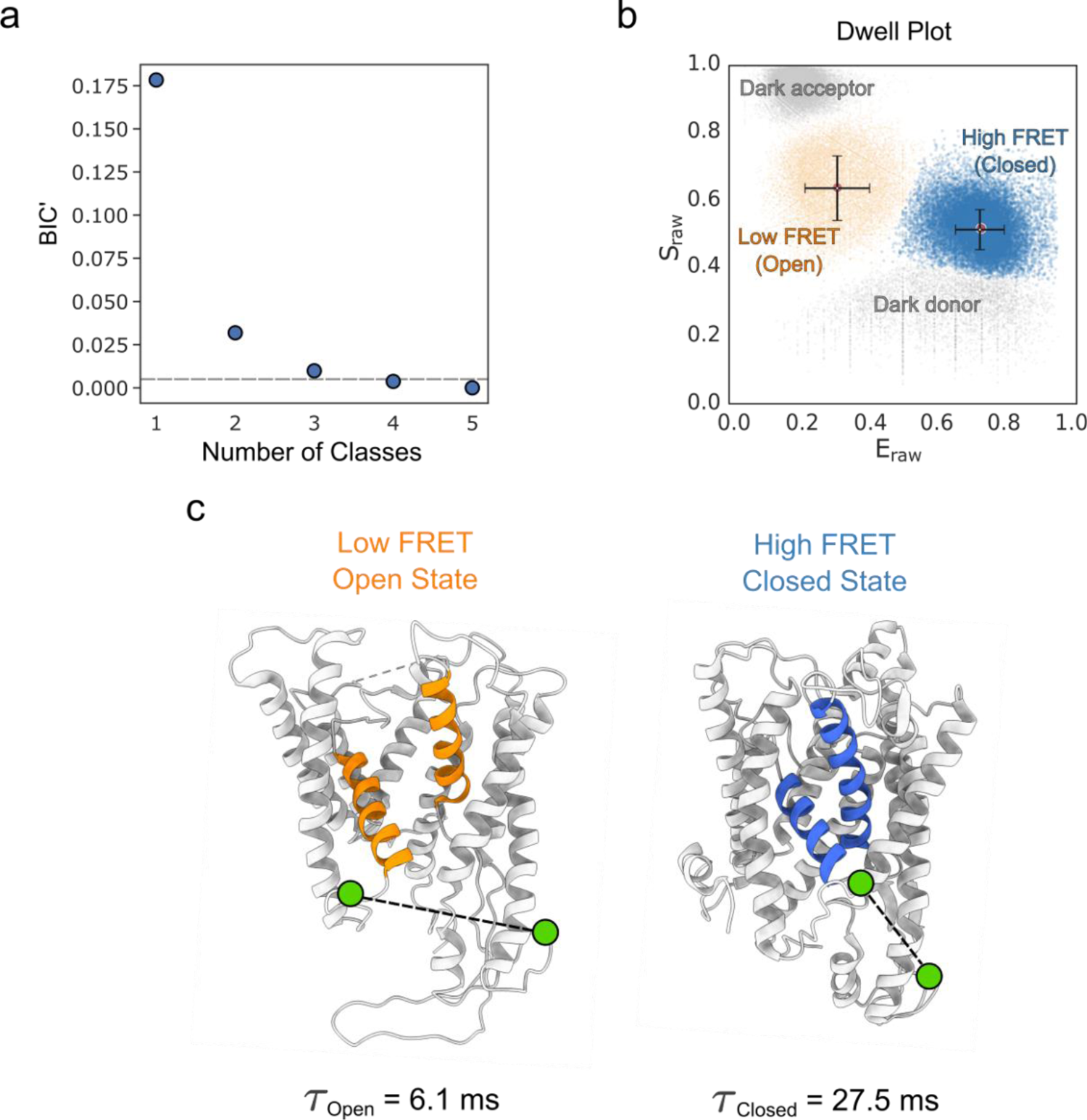
Conformation and dynamics of SecYEG protein channel and the adjacent LG are discernible by smFRET on the millisecond timescale. a) The modified Bayes Information Criterion (BIC’) derived by mpH^2^MM analysis of SecYEG alone in liposomes. The different state-models indicate that four classes best describe the data (*i.e.* BIC’ < 0.005, grey dashed line). b) Scatterplot of dwells for SecYEG alone in liposomes showing the dwell E_raw_ and S_raw_ derived from the mpH^2^MM analysis. The black bars represent the E_raw_ and S_raw_ standard deviations for each respective population. See Supplementary Fig. 4 for more detailed raw data. c) Models highlighting the SecY LG in the open (low FRET) and closed (high FRET) conformations (PDBs: Open=3DIN^30^, Closed=1RHZ^18^). The two fluorophore labelling positions (A103C and V353C on SecY, shown as green circles and connected with a black dashed line) in the open and closed state are indicated. The dwell time (τ_open_ & τ_closed_) in each state in SecYEG alone is shown below each Fig. part.

The dynamic nature of the equilibrium between the closed and open states is manifested as transitions on the millisecond time scale (detected by mpH^2^MM analysis, Fig. 2c and Supplementary Methods) and confirmed by burst variance analysis (Supplementary Methods and Supplementary Fig. 4e)). Hence, even in the absence of SecA or translocating protein, the SecYEG core-complex is conformationally dynamic, switching between open and closed states on the millisecond timescale.

Previous studies indicated that the conformation of SecYEG is controlled by the nucleotide occupancy of SecA^8,10, 34, 39^. The complex is also known to respond to the presence of a translocating pre-protein substrate^8,19, 34, 39^, or to the addition of an isolated signal sequence peptide (SS)^39–41^. We therefore collected single-molecule FRET data under a range of different conditions (± SecA, various nucleotides and a SS peptide) for mpH^2^MM analysis. In each case, the data (Supplementary Fig. 4-11) are best described by the four classes (Fig. 3a) assigned to the two conformational states (open and closed) and the irrelevant photophysical donor only and acceptor only events (Fig. 3b). While the two states of interest were always present, their relative proportions changed considerably. For SecYEG alone, there is observable rapid interchange on a millisecond timescale between the low and high FRET states (burst variance analysis (BVA)^42^; Supplementary Fig. 4e-11e). Thus, using channel opening as a reaction coordinate, the state of the SecYEG:A complex cannot be described by static structures in slow exchange. Instead, the behaviour is better described by a dynamic equilibrium between open and closed states that interconvert on the millisecond timescale.

### SecA and its associated nucleotide modulate the dynamics of the protein channel within SecYEG

The mpH^2^MM methodology allows extraction of the dwell times of the two states of SecYEG (τ_open_ and τ_closed_). These serve as dynamic equilibrium descriptors, and can be visualised in two-dimensional plots (Fig. 4a,b). Here, changes in the time spent in the open or closed states can be readily visualised (Fig. 4a,b (diagonal lines); Fig. 4c). The addition of SS to SecYEG (in the absence of SecA), which becomes wedged into the open lateral gate^17, 32, 41^ (Fig. 1), causes ‘unlocking’ of the LG, priming SecYEG for protein transport^43^. Within the dynamic equilibrium context, SS binding dramatically increases the total time SecYEG spends in the open state, not by increasing τ_open_, but by decreasing τ_closed_ (from 27.5 ± 4.6 ms to 1.3 ± 0.6 ms) (Fig. 4a). This equates to a ∼ 4-fold increase in the open state percentage (Fig. 4c, Supplementary Table 2). On the other hand, the addition of SecA (in the absence of nucleotide or SS) tips the balance towards the closed state ∼ 2-fold, by decreasing τ_open_, with little effect on τ_closed_ (Figs. 4a,c).

**Fig. 4.**
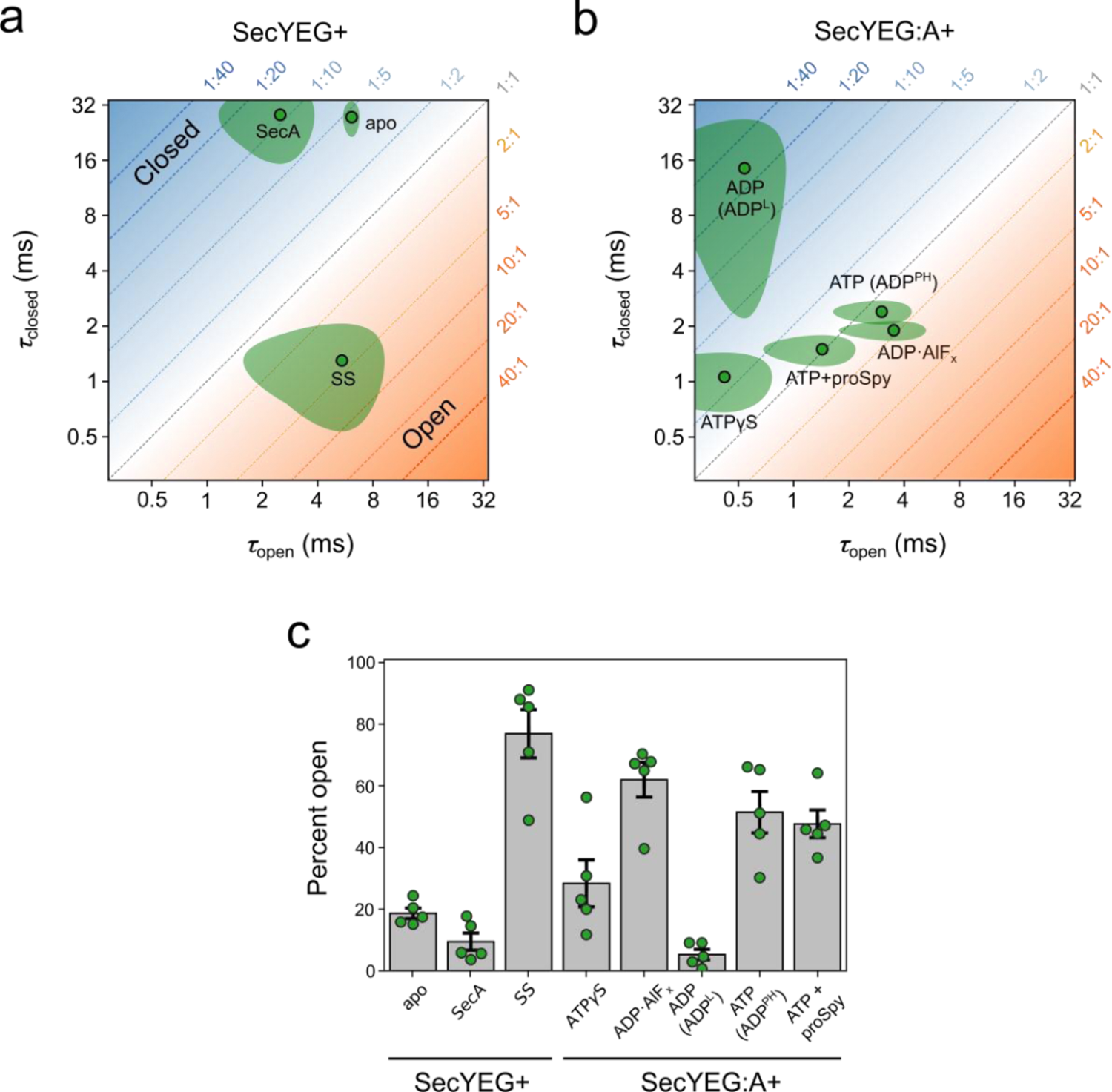
Dynamics of the SecYEG protein channel and LG. a) SecYEG channel dwell times in the open (τ_open_) and closed (τ_closed_) states for SecYEG with the addition of (SecYEG+): nothing (apo), SecA or SS. The 90% confidence areas are shaded green with the average dwell times shown as green points with black outline. The ratio of time spent in each conformational state is shown as a slanted dotted line with the associated fraction in each state (open:closed) in orange/ blue in the plot margins. b) As in panel a), but with the addition of various nucleotides and pre-protein to the SecYEG:A complex (SecYEG:A+). c) The percent open (i.e. τ_open_/(τ_open_+τ_closed_)x100) for data shown in panels a) and b). Error is given as 90% confidence intervals. Raw data for dwell times and percent open are shown in Supplementary Tables 1 and 3. smFRET raw data are shown in Supplementary Fig. 4-11.

Channel dynamics within the SecYEG:A complex respond to the nucleotide bound to SecA, even in the absence of a pre-protein substrate (Fig. 4b,c). In the presence of SecA there are three distinct types of dynamic behaviour: (i) mostly closed (95 ± 4 %) and relatively static in the SecYEG:A:ADP complex; τ_open_ is short (0.5 ± 0.3 ms) compared with the long-lived τ_closed_ (14.5 ± 9.4 ms)); (ii) highly dynamic, but still biased towards closed (71 ± 16 %) seen in the SecYEG:A:ATPγS complex (τ_open_ is unaffected compared with SecYEG:A:ADP (0.4 ± 0.3 ms) but τ_closed_ is dramatically decreased (1.1 ± 0.3 ms)); and (iii) moderately dynamic with a larger time spent in the open state (62 ± 12 %) found in the SecYEG:A:ADP·AlF_x_ complex, which is thought to resemble a post-hydrolysis transition state^44, 45^ (τ_open_ is increased (3.5 ± 1.3 ms) and τ_closed_ is marginally affected (1.9 ± 0.2 ms)).

The rate limiting step in the SecA ATPase cycle is ADP release^28^. Hence, in the absence of pre-protein, the complex spends most of its time (∼ 99%) bound to ADP. We thus expected channel dynamics in the SecYEG:A:ATP steady-state complex (formed by including 1 mM ATP in the experiment (K_M[ATP]_ = 51.1 ± 7.8 µM^28^)) to resemble the dynamic position displayed by the SecYEG:A:ADP complex. Remarkably, however, the dynamic equilibrium position for the SecYEG:A:ATP complex resembles closely that of the post-hydrolysis state SecYEG:A:ADP·AlF_x_, and is significantly different to the SecYEG:A:ADP and SecYEG:A:ATPγS complexes (Fig. 4b). This tells us that ATP must already have been hydrolysed (otherwise the complex would exhibit dynamics similar to the ATPγS state) and P_i_ released (P_i_ release is very fast; 11.5 ± 0.07 s^-1^ compared with a k_cat_ of 0.15 ± 0.02 s^-1^ ^28^). Therefore, SecA must be in a particular ADP bound state, which can only be achieved through hydrolysis of ATP. We designate this state as the post-hydrolysis (PH) form (named SecYEG:A:ADP^PH^) to distinguish it from the predominantly closed complex with loosely (L) associated ADP (SecYEG:A:ADP^L^), formed by the simple addition of ADP to SecYEG:A. This newly assigned state of SecA clearly imparts distinct dynamics to the SecYEG channel and hence is relevant to the functional cycle of the SecYEG:A complex.

### Impact of a pre-protein client on the conformation of the protein channel

We next examined how the dynamic equilibrium between the open and closed states responds to the translocation of a pre-protein (spheroplast protein Y; proSpy). Spy is a 15.9kDa periplasmic chaperone^46, 47^ which has been used previously as a model protein for translocation through SecYEG^29, 48, 49^. The rate of ATP hydrolysis by SecA during translocation of proSpy increases > 40-fold (one ATP hydrolysed every 0.15 s compared with one ATP every 6.66 s in the absence of proSpy (Supplementary Table 3)). This increase in the observed rate of ATP hydrolysis in the presence of pre-protein is caused by an increased rate of ADP release, increasing the fraction of time spent by the complex in the ATP-bound pre-hydrolysis state from an average of ∼1% to ∼37%^28^. Consistent with this, the steady state dynamic equilibrium of the SecYEG:A:ATP+proSpy complex no longer coincides with the post-hydrolysis ADP state (Fig. 4b; SecYEG:A:ADP^PH^). Instead, it maps to an area between SecYEG:A:ADP^PH^ and the pre-hydrolysis SecYEG:A:ATPγS state (Fig. 4b); thus approximating a steady-state mixture of pre- and post-ATP hydrolysis states.

Taken together, the smFRET data show that the slow ATPase cycle of SecA which occurs on a > 100 millisecond timescale^10, 28^ modulates intrinsically fast (millisecond timescale) dynamics of the SecYEG core-complex. A distinct, post-hydrolysis, post-phosphate release ADP-bound state (ADP^PH^), which is dynamic (τ_open_ = 2.8 ± 1.3 ms and τ_closed_ = 2.4 ± 0.4 ms) and has an open to closed ratio of ∼1:1 (51 ± 14% open), dominates the steady state ATPase cycle in the absence of pre-protein. Thus, the rate limiting step is a conformational change that involves the SecYEG channel and adjacent LG and is associated with the conversion of SecYEG:A:ADP^PH^ to a state from which ADP is readily released, most likely the equilibrium SecYEG:A:ADP^L^ state. During translocation this conformational change is accelerated, allowing for faster ADP release and therefore ATPase turnover (>40-fold, Supplementary Table 3).

### SecYEG channel and lateral gate dynamics are linked to the rate of protein translocation

Next, we explored the role of the rapid millisecond SecYEG dynamics, and its control by the ATPase cycle of SecA, for protein transport. To do so we utilised the widely studied variant of SecYEG PrlA4, which contains two amino acid substitutions in SecY, F286Y and I408N (in TM7 & TM10 respectively (Supplementary Fig. 3))^50, 51^. This variant was produced by an *E. coli* strain selected to suppress the effects of a defective signal sequence. This suppression is achieved by the resulting SecYEG-PrlA4 complex being primed in an ‘unlocked’ conformation, which would otherwise require the docking of a functional SS at the LG^18, 51–54^. Given that SS insertion dramatically affects channel dynamics (Fig. 4a), we hypothesised that the PrlA4 variant may also exhibit perturbed dynamics such that the open state is promoted.

Experiments measuring the rate of translocation of proSpy (see Methods) showed that the PrlA4 variant has a translocation rate ∼3.4-fold greater (9.70 ± 0.02 aa.s^-1^) than that of wild-type SecYEG (2.85 ± 0.03 aa.s^-1^) (Fig. 5a, Supplementary Table 4 and 5), while having approximately the same ATP hydrolysis rate by SecA during translocation (Fig. 5b, Supplementary Table 3). These data reveal two important points: (i) the SecYEG-PrlA4 variant is more efficient at translocation of pre-protein substrates (Fig. 5c, 1.38 ± 0.09 aa/ATP compared to wild-type SecYEG 0.44 ± 0.03 aa/ATP) and (ii) the ATP hydrolysis cycle seems not to be strictly correlated with the rate of proSpy translocation, which would be expected if SecA was acting alone as a stepping motor.

**Fig. 5.**
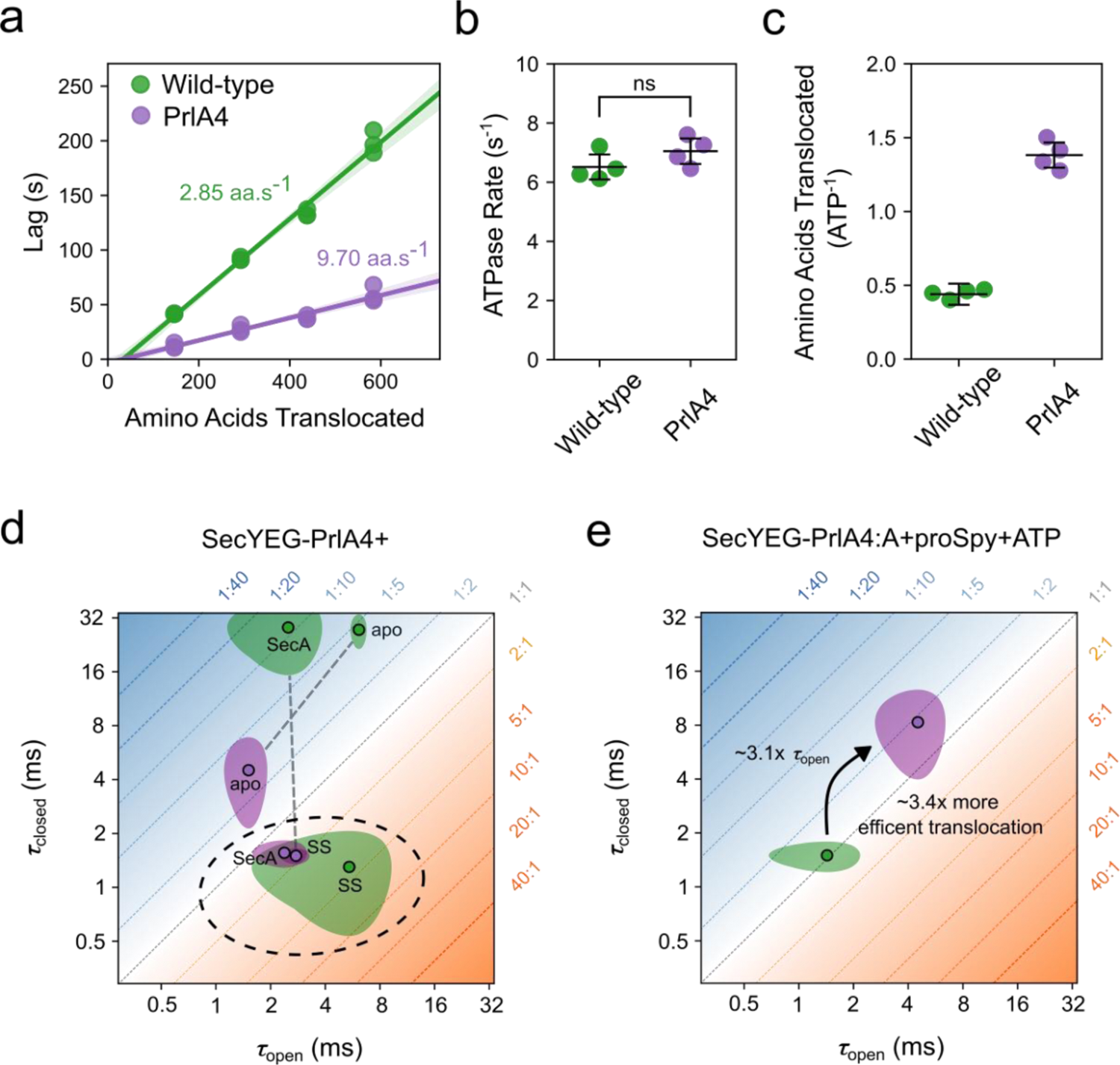
The PrlA4 variant is a more efficient translocase than the native SecYEG complex. a) Rates of transport of proSpy for SecYEG and SecYEG-PrlA4 as measured by the NanoLuc split-luciferase assay. The assay measures the time lag before the onset of chemiluminescence due to binding of high affinity β-strand, which is inserted at various positions into a 4x tandem repeat of proSpy, to NanoLuc luciferase fragment encapsulated within SecYEG proteoliposomes (Methods and Supplementary Methods, data shown in Supplementary Table 6 and 7). A linear regression fit is shown as a solid line, with the 95% confidence interval shown (95%CI) shaded in the respective colour. b) Steady-state rates of ATP hydrolysis by SecA translocating proSpy through SecYEG vs SecYEG-PrlA4 (ns, p-value = 0.18). c) The number of amino acids translocated by SecYEG per ATP hydrolysed by SecA during transport of proSpy in wild-type (0.44 ± 0.03 (95%CI)) and PrlA4 (1.38 ± 0.09 (95%CI)). d) LG dwell times (shown as 90% confidence interval plots) of SecYEG-PrlA4, SecYEG-PrlA4:A and SecYEG-PrlA4:SS (purple) compared with the wild-type complex (green, data for the latter are reproduced from Fig. 4a for comparison). Conditions representing the ‘unlocked’ state of the LG (*i.e.* primed for transport and have an equilibrium position favouring the open state) are circled with a grey dashed line. e) LG dwell times (shown as 90% confidence interval plots) of the translocation of proSpy, coloured as in (d) with the steady-state dwell time in the open state and translocation efficiency compared with that of the wild-type.

To determine how the amino acid substitutions in the PrlA4 variant affect channel dynamics, we utilised the same labelling as for wild-type SecYEG at residues A103C and V353C (see Methods) and again analysed the dynamics of the channel using by smFRET. The results revealed that PrlA4 exhibits the same two FRET states observed in wild-type SecYEG (Supplementary Fig. 12-15), corresponding to the same open and closed conformations. However, the interconversion between these states is drastically different (Supplementary Table 6). In the apo complex (in the absence of SecA/ nucleotides/ SS) PrlA4 is more dynamic than SecYEG (τ_open_ = 1.5 ± 0.3 ms and τ_closed_ = 4.5 ± 1.8 ms (Fig. 5d), compared with τ_open_ = 6.1 ± 0.5 ms and τ_closed_ = 27.5 ± 4.6 ms for wild-type SecYEG (Fig. 4a)). While more mobile, the overall percentage of time spent in the open state is roughly the same (PrlA4 = 26 ± 4 % compared to wild-type SecYEG = 19 ± 4 %, Supplementary Table 2 and 7).

Upon the addition of SecA, the dynamic equilibrium of PrlA4 is shifted significantly towards the open configuration (60 ± 6 %), with a rate of dynamic interchange similar to that observed for wild-type SecYEG bound to the SS peptide (τ_open_ = 2.4 ± 0.6 ms and τ_closed_ = 1.6 ± 0.2 ms) (Fig. 5d). Therefore, PrlA4 can be unlocked by SecA (more open) without the need for a SS, explaining its ability to translocate pre-proteins with defective signal-sequences. Previous reports suggested that the PrlA4 variant cause a general ‘relaxation’ in SecYEG, rather than a specific conformational change which allows for the bypassing of signal sequence recognition^50^. Our findings support this notion, but additionally show that association of SecA is necessary and sufficient for the complex to adopt the unlocked state.

During translocation, SecYEG-PrlA4:A:ATP+proSpy alternates between long dwells in both the open and closed channel states (τ_open_ = 4.5 ± 1.5 ms τ_closed_ = 8.3 ± 3.3 ms), in contrast to the wild-type complex (SecYEG:A:ATP+proSpy), which is more dynamic under the same conditions (τ_open_ = 1.4 ± 0.6 ms and τ_closed_ = 1.4 ± 0.5 ms) (Fig. 5e). Interestingly, the total percentage of time spent in the open state by the PrlA4 during translocation (36 ± 10 %) is similar to that of wild-type SecYEG (48 ± 9 %) (Supplementary Tables 2 and 7). Hence, the equilibrium position between open and closed states is not responsible for the phenotype of *prlA4,* but rather the time the channel spends in the open state is perturbed, with the complex residing in the open state ∼3.1-times longer than in the wild-type complex.

Given that passive polypeptide diffusion has been shown to contribute significantly to the kinetics of translocation^9^, the extended τ_open_ would allow a longer time for free diffusion of the pre-protein through the SecY pore; thereby, increasing the probability that rate-limiting charged and bulky residues could pass through the pore before closure^48^. Consistent with this, the increase in the dwell time of the open conformation in PrlA4 corresponds to the observed increase in protein translocation rate and efficiency (∼3.4-fold; Fig. 5e). Together, the results show that the altered translocation properties of the PrlA4 variant are not due to SecA motor action, but instead correlate with the duration of the open state of the protein channel in SecYEG.

## Discussion

### Energy landscape steering of SecYEG protein channel dynamics

In this study we have exploited the powers of smFRET analyses to show that rapid millisecond dynamic interchange between two conformational states is an intrinsic property of SecYEG, independent of SecA. The closed state is the most highly populated (∼81%) for SecYEG alone at equilibrium. Binding of SecA (without nucleotide) to SecYEG shifts the equilibrium to further promote channel closure (to > 91%) by decreasing the dwell time of the open state. This allosteric effect is further modulated by the nucleotide state of SecA.

Based on previous biophysical studies from our own group and elsewhere^8–10, 23, 29, 32–34, 39, 43, 48^ it was anticipated that the conformations of the SecYEG protein channel would mirror the slow (>100 ms) ATPase cycle of the associated SecA. In other words, discrete states of the channel would exist that are strictly constrained by the bound nucleotide on SecA–interconverting upon ATP hydrolysis and ADP exchange. As we show here, it is only in the SecYEG:A:ADP^L^ complex that the channel may be considered quasi-static (∼95% closed), and that the binding of ATP, or non-hydrolysable analogues, to SecA shifts the population of the dynamic equilibrium towards the open state. This is achieved by inducing fast, millisecond time scale interconversion between the open and closed states. The SecA nucleotide dependent control over rapid SecYEG channel and LG dynamics can be visualised with the help of simplified energy landscapes projected onto a reaction coordinate of opening and closure (Fig. 6a). Within this framework SecA ‘steers’ the energy landscape by changing the relative depth of the two minima (equilibrium) and the barrier height between them (interconversion rate).

**Fig. 6.**
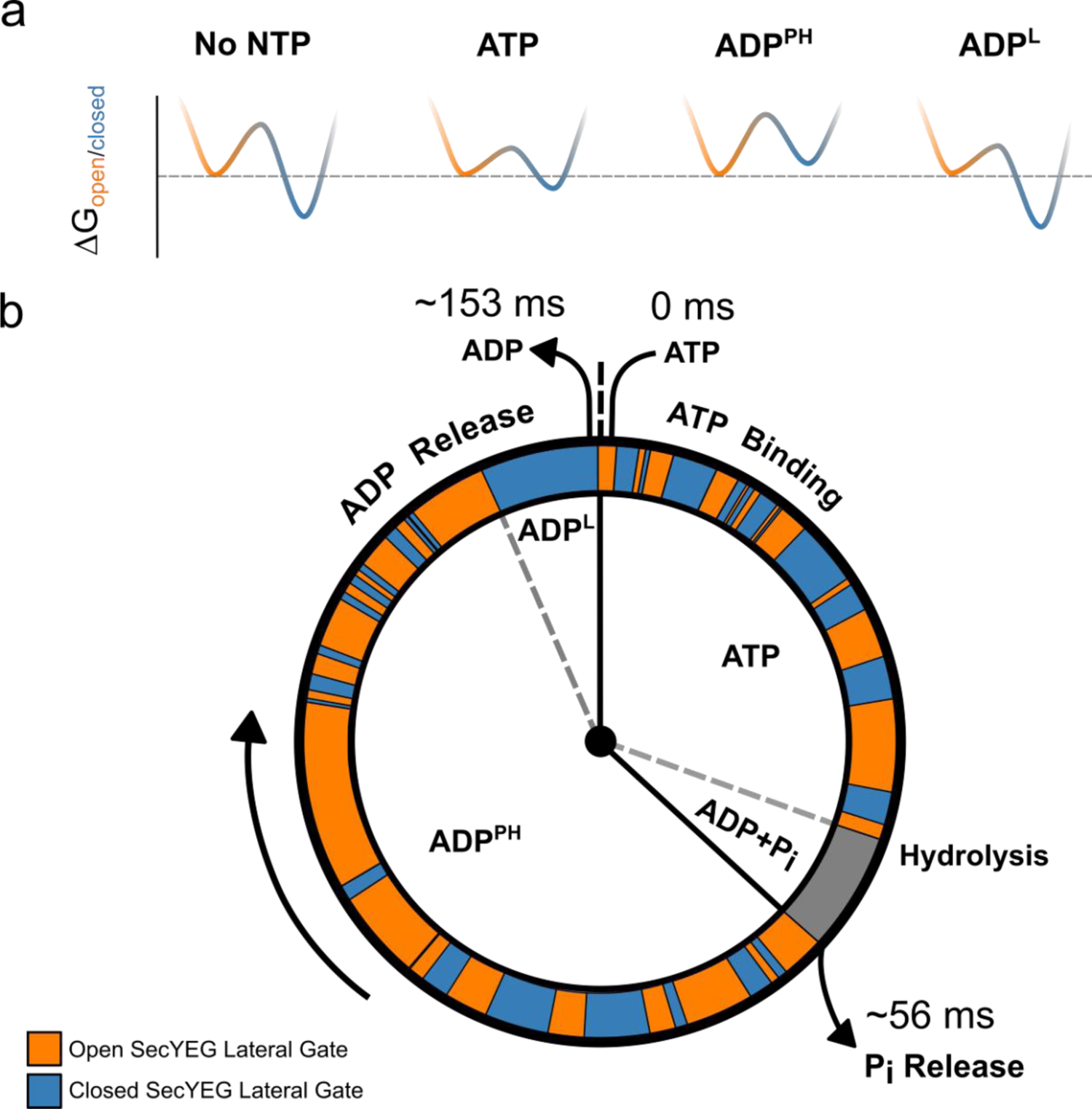
Observed SecYEG protein channel dynamics as a function of the ATP hydrolysis cycle of SecA. a) Free energies between conformational states of the SecYEG were estimated from data for the following states: No NTP = SecYEG:A, ATP = SecYEG:A:ATPγS, ADP^PH^ = SecYEG:A:ATP, ADP^L^ = SecYEG:A:ADP. The closed state well is coloured blue while the open state is orange. b) The ATP hydrolysis cycle of SecA is shown as a ‘clock’ with a simulated trajectory of channel conformations shown in orange (open) and blue (closed) around the perimeter. The representative trajectory was simulated with the experimentally derived rates of interconversion. The time spent in each step during translocation is calculated from the k_cat_ of ATP hydrolysis (6.52 ± 0.42 s^-1^) (Fig. 5b) and the time taken for phosphate release^28^ (17.9 ± 0.15 s^-1^).

To better visualise the relationship between the hydrolytic cycle of SecA and the dynamics of the SecYEG channel, we projected a representative interconversion trajectory of the open and closed states onto an ‘average’ ATPase cycle (during translocation; Fig. 6b). The time SecA spends associated with each nucleotide was determined using two measured parameters: i) the steady-state ATPase rate (Fig. 5b), and ii) the rate of phosphate release^28^. We further assumed that ATP binding at the millimolar concentration used is much faster than any subsequent steps^28, 55^, and thus the apo SecYEG:A (without nucleotide) does not contribute to the cycle. Likewise, since phosphate release is fast^28^, we assume the system spends a relatively short time in the ADP+P_i_ post-hydrolysis state. Currently, we do not know if the transient pre-phosphate release step is more like the ATP or ADP^PH^ bound complexes, or something in between. During translocation, the steady-state mixture of nucleotide associated SecYEG:A complexes can be best approximated by a mixture of the ATP-bound (ATPγS) and ADP^PH^ (ATP) forms. Hence, we estimated that the ADP^PH^ form of the complex to dominate the ADP associated phase of the cycle, with the contribution of the ADP^L^ (loosely associated) form being marginal.

The kinetic cycle we describe highlights the stark difference in timescales between SecYEG channel dynamics and those of ATP hydrolysis by SecA. The results clearly demonstrate that SecYEG and SecA cannot be directly coupled, and that thermal fluctuations of the channel and the adjacent LG play an important role in the pre-protein translocation mechanism. These data support previously proposed models suggesting a dynamic equilibrium between open and closed conformations of SecYEG based on lateral gating being strongly dependent on temperature^56^. Thus, the functional state of the SecYEG:A complex cannot be represented by a single structure. Instead, protein translocation will involve an ensemble of co-existing conformations interchanging on a rapid timescale; effectively in a dynamic equilibrium controlled by the SecA ATPase cycle. Thereby, SecA ‘steers’ the conformational ensemble through a sequence of different dynamic equilibria by modulating the landscape according to its nucleotide state (Fig. 6).

### Fast protein channel and LG dynamics are integral to the mechanism of translocation

Previous cross-linking experiments demonstrated that channel and LG opening is essential for translocation^19^ and that closure slows down ADP release^8^. In contrast, translocation models based solely on SecA motor action predict the translocation speed to be directly linked to the ATP hydrolysis rate, independent of channel dynamics^10, 33^. We found that the SecY-PrlA4 variant exhibits a faster (∼3.4 times faster) pre-protein translocation rate at the same ATP cost. smFRET analysis showed unequivocally that the dynamics of the SecYEG channel is an integral part of the translocation mechanism. Importantly, we determine that the site-specific changes in PrlA4 do not affect the equilibrium position of open versus closed channel, but significantly increase the dwell times of both states. This increase in dwell time (∼3.1 times slower interconversion) correlates well with the increased rate of translocation (∼3.4 times faster). The longer dwell time of the open state would increase the probability of charged and bulky residues within the pre-protein sequence being translocated through the channel. This observation might also explain the reported ability of the PrlA4 variant to translocate partially folded pre-protein substrates^50^.

Our data, alongside recent single-molecule studies of SecA^10, 57^, indicate that SecYEG and SecA operate on distinct, but stochastically coupled timescales that are not compatible with a directly coupled power stroke. The evolution of such a mechanism with interacting, but dynamically independent, components could be versatile and widespread. The shallow energy landscape of the different states of the core SecYEG complex enables rapid access to different conformational states and interconversion between them. The ‘steering’ of these energy landscapes, in conjunction with accessory factors, could then be exploited for different activities, such as insertion of trans-membrane helices into the bilayer.

Such a dynamic allosteric mechanism is not unique to the SecYEG translocon, since similar nucleotide-dependent steering of rapid dynamics has also been recently shown for adenylate kinase^58^ and the AAA+ ring ATPase, ClpB^59^, which were also revealed by advances in single-molecule fluorescence techniques. Other ATP-driven pumps also have proposed mechanisms incorporating a series of static states associated with or without different nucleotides (*e.g.* the ABC transporters^60^). In view of the work described here it would not be surprising if rapid conformational interconversion between key transport states, *i.e.* inward and outward facing, are also subject to the influence of the various stages of the ATP hydrolytic cycle, operating at a different timescale. It may indeed be the case that dynamic allostery involving motions occurring across multiple time scales coupled *via* energy landscape steering may be ubiquitous among complex molecular machines.

## Methods

### Protein Expression, Purification and Labelling

Protein production, purification and labelling were performed according to protocols published previously^8,29, 61, 62^. *E. coli* SecYEG with two unique Cysteine (Cys, C) residues, A103C and V353C, in SecY (SecY_A103C-V353C_EG) was used and labelled with ATTO 565 as a donor dye and ATTO 643 as an acceptor dye for single-molecule FRET measurements, as previously described^8^. The prlA4 variant^50, 53^, which also contained the mutations A103C and V353C, was created by site-directed mutagenesis before being purified and labelled as described for its wild-type equivalent. Note that under the conditions used (100 µM SecY_A103C-V353C_EG, 200 µM each of ATTO 565 maleimide and ATTO 643 maleimide), labelling of each Cys with each fluorophore is random. The translocation precursor, proSpy, which contains Spy with its natural signal sequence, was expressed in an *E. coli* strain defective in the export of secreted proteins and purified to homogeneity as described by Pereira et al.^49^. Full details can be found in the Supplementary Methods.

### ATPase Assays

ATPase assays were performed and analysed as described by Gold et al. ^62^,modified to allow data collection on a BioTek Synergy Neo2 plate reader; full details can be found in the Supplementary Methods.

### Transport Assays

Transport assays were performed as described by Pereira et al.^49^, and the data analysed as in Allen et al.^29^. Full details can be found in the Supplementary Methods.

### Single-molecule Sample Preparation

Proteoliposomes were prepared as in Allen et al.^8^ with minor adjustments. Labelled SecY_A103C-V353C_EG was reconstituted to a concentration of 37.5 nM into *E. coli* polar lipid extract (10060C, Avanti Polar Lipids) at a volume of 400 μl and a concentration of 5 mg/ml in TKM buffer (20 mM Tris, 50 mM KCl, 2 mM MgCl_2_, pH 7.5). The mixture was then extruded to form proteoliposomes with a diameter of 100 nm; at this concentration and size, most liposomes are expected to contain either 0 or 1 copy of SecY_A103C-V353C_EG^63^. The extrusion step was performed on a heating block set to 40 °C (610000-1EA, Avanti Polar Lipids). The resulting mixture was dialysed overnight using D-Tube Dialyzer Mini with a Molecular weight cut-off of 12-14 kDa (Sigma Aldrich) in TKM buffer before being stored at 4 °C.

### Single-Molecule Data Acquisition

Single-molecule Förster resonance energy transfer (smFRET) experiments were performed on a custom-built confocal epi-illuminated microscope (Supplementary Methods) in a standard inverted-stage configuration with a pulsed interleaved excitation regime^38^. Samples were measured in an 8 well sample chamber (80827, Ibidi) which had been coated in bovine serum albumin (BSA) to prevent any adhesion of the sample to the chamber. Coating of the sample chamber was achieved by pipetting 500 μl of 1 mg/ml BSA solution which had been filtered through a 0.22 μm membrane and leaving at room temperature for 20 minutes. After this time, the BSA solution was removed and the sample chamber was rinsed thoroughly with MilliQ water before being left to air dry. Samples were measured at a concentration of 30 pM in TKM buffer supplemented with 1 mM aged Trolox to help reduce blinking and photobleaching of the fluorophores. Trolox was aged to form a fraction of oxidised Trolox (Trolox-qinone) which then acts with Trolox according to a reducing and oxidizing system scheme^64, 65^. Aged Trolox was made by adding 1 mM of Trolox to TKM buffer and left overnight on a shaker at 4°C to dissolve before being filtered using a 0.22 μm membrane. The chamber was covered during data acquisition to help prevent evaporation of the sample. Relevant components for each condition were added immediately before measurement to final concentrations of 1 μM SecA, 1 μM SS, 0.7 μM proSpy and 1 mM ATP, ATPγS, ADP and ADP·AlF_x_. ATP depletion was negligible due to low SecYEG:A concentration and low intrinsic turnover of free SecA^28^. Each condition was measured 5 times for 1 hour (apart from ATP+proSpy, where each measurement for 20 minutes) with fresh proteoliposome preparations. Supplementary Fig. 16 shows the variability in the data of the five fresh proteoliposome preparations for SecYEG apo.

### Single-Molecule Data Analysis

Data were analysed using the FRETBursts python package^66^. A burst search with a minimum threshold of 6x the background signal in the donor and acceptor channels respectively and a minimum burst size of 50 photons was used to distinguish single molecule events. Further details on burst selection, mpH^2^MM and statistical analyses are given in the Supplementary Methods. Due to the random nature of insertion of SecY_A103C-V353C_EG into the liposomes, ∼50% of the protein was oriented in such a way that the SecA binding interface was inaccessible^8,13, 67^, therefore we applied a correction factor to account for the remaining 50% apo (Supplementary Methods).

## Acknowledgments

This work was funded by the BBSRC: J.A.C., R.T. and S.E.R. (BB/T008059/1); W.J.A. and I.C. (BB/V001531/1); D.W.W and I.C. (BB/T006889/1). SER holds a Royal Society Professorial Research Fellowship (RSRP\R1\211057). J.A.C., T.F. and R.T. are supported by the European Regional Development Fund-Project (CZ.02.1.01/0.0/0.0/15_003/0000441) and the Czech Science Foundation (20-11563Y). J.A.C. and T.S. also acknowledge Leeds Beckett University for funding. We thank David Brockwell for his feedback and discussion and G. Nasir Khan for his excellent technical support. We also thank Matthew Watson for performing data collection in the early stages of the project.

## Author Contributions

J.A.C. and T.F. built the instrumentation for single-molecule experiments, prepared the samples, acquired the data and performed the data analysis. D.W.W. and W.J.A. performed protein expression, purification, labelling and characterisation. I.C., R.T., S.E.R., T.F. and T.S. acquired funding and supervised the project. All authors contributed to planning of experiments, interpretation of results and writing of the manuscript.

## Data Availability

The data that support this study will be made freely available from the corresponding authors upon request. All ATPase assay, transport assay, single-molecule FRET raw data files and scripts for analysis will be made available via the University of Leeds Research Data Repository.

