## Supplementary Information for "Energy landscape steering mediates dynamic coupling in ATP-driven protein translocation by the bacterial Sec machinery"

#### Supplementary Methods

##### Protein expression, purification, labelling and reconstitution

All SecYEG variants, cloned in pBAD/*myc*-His C and with an N-terminal 6His tag on SecE, were taken from our laboratory collection. Plasmids were transformed into competent *E. coli* C43 cells. A 100 ml pre-culture of 2xYT broth containing 100 µg.ml$^{-1}$ ampicillin was inoculated and incubated overnight at 200 RPM and 37°C. The following morning, Flasks containing 2xYT with 100 µg.ml$^{-1}$ Ampicillin were inoculated with pre-culture (15 ml per 2 L, in 5 L flasks) and incubated at 200 RPM and 37°C. On reaching an $A_{600}$ of 0.8, expression was induced by addition of arabinose to 0.2 % (w/v). After 3 hours, cells were harvested by centrifugation (5000 g, 15 mins) and resuspended in 50 ml 20 mM Tris pH 8, 300 mM NaCl, 10 % glycerol (TSG300). The cells were passed twice through a cell disruptor (Constant Systems Ltd.) at 25 kPSI and membranes clarified by centrifugation (160,000 g, 45 mins, 4°C). The membrane pellets were then washed gently in TSG300 and stored at -20˚C until purification.

For purification, membrane pellets were resuspended to 42.5 ml in TSG and homogenised with a potter. A 10 % n-Dodecyl β-D-maltoside (DDM, Glycon) stock was added to 1 % final, and membranes were incubated gently rotating at 4°C for 30 mins. Insoluble material was removed by centrifugation (160,000 g, 45 mins, 4°C). A 25 ml bed volume nickel immobilised metal affinity chromatography column linked to an Akta FPLC (GE Healthcare) was equilibrated in TSG300 + 0.1 % DDM. The solubilised membranes were loaded onto the column, which was subsequently washed with TSG300 + 0.1 % DDM + 30 mM imidazole until the A280 stabilised. On completion, bound material was eluted with TSG300 + 0.1 % DDM + 330 mM imidazole. The eluate was subsequently loaded onto a HiLoad 26/600 Superdex 200 pg size exclusion column with a 25 ml bed volume Q sepharose fast flow (GE healthcare) column connected in tandem, pre-equilibrated in TSG130 (as TSG300, but with only 130 mM NaCl) + 0.02 % DDM. Purified SecYEG was concentrated to approximately 200 µM in a 50 kDa cut-off centrifugal concentrator (Sartorius Stedim) and stored at -80°C for future use.

Before labelling, $SecY_{A103C-V353C}EG$ was treated with 100 mM DTT for 30 mins to fully reduce all cysteines. The sample was then passed over a 20 ml Superdex 200 10/300 GL size exclusion column equilibrated in TSG130 + 0.02 % DDM (to remove excess DTT), and the protein peak concentrated to ~100 µM, then immediately labelled with 200 µM ATTO 565 maleimide and 200 µM ATTO 655-maleimide, in the dark at 4˚C for 2 h. The reaction was quenched by addition of 10 mM DTT, and excess probe removed by size exclusion exactly as for the DTT. The resulting protein was concentrated and stored at -80˚C until use.

We employed exactly the same translocation substrate constructs as in Allen et al.[1]: the soluble periplasmic chaperone proSpy followed by three tandem repeats of the Spy mature domain, with each Spy sequence followed by pep86 (to allow detection by NanoLuc; $pSpy_{4x}$).

For each substrate, three copies of pep86 were scrambled such that they have the same amino acid composition but no longer interact with 11S, leaving only one functional pep86 sequence. The resulting proteins are identical except for the number of amino acids that must be transported before the functional pep86 reaches the vesicle lumen and interacts with 11S. Expression plasmids for these constructs (in pBAD/*myc*-His C) were transformed into *E. coli* MM52 cells. A 250 ml 2xYT preculture containing 100 µg/ml ampicillin was grown overnight at 30°C and 200 RPM in a 2L flask. 1 L 2xYT containing 100 µg/ml ampicillin, prewarmed to 39°C, was added to the overnight culture before incubation at 39°C and 200 RPM. After 45 minutes, expression was induced with addition of arabinose to 0.2%. 3 hours later cells were harvested, resuspended and lysed as for SecYEG (described above), but with 50 ml resuspension buffer of 50 mM Tris-HCl pH 8.0 and 500 mM NaCl (TS500 buffer). The sample was clarified by centrifugation (20,000 g, 30 minutes, 4°C) and the soluble fraction taken. Nickel affinity purification was conducted as for SecYEG, but columns were washed in TS500 containing 30 (wash) or 330 mM (elution) imidazole. The eluate was passed over a Q-sepharose column to remove any remaining contaminants, and the flowthrough exchanged into 20 mM Tris pH 8 and 6 M urea and concentrated to ~200 µl final volume with a 10 kDa centrifugal concentrator and stored at -80°C for future use.

$_{His6}$SecA cloned in pTrc99a from our laboratory collection was transformed into competent *E. coli* BL21(DE3), used to inoculate a 100 ml culture of LB broth containing 50 µg.ml$^{-1}$ kanamycin and incubated overnight at 200 RPM and 37°C. The following morning LB supplemented with 50 µg.ml$^{-1}$ kanamycin was inoculated with pre-culture (15 ml per 2 L, in 5 L flasks), left to incubate at 200 RPM and 37°C until the $A_{600}$ reached 0.6, then expression was induced with 1 mM IPTG for 1.5 hours. Cells were harvested by centrifugation (5000 g, 15 mins), resuspended in 100 ml TKM (20 mM Tris pH 8, 50 mM KCl, 2 mM $MgCl_2$), lysed as described for SecYEG and clarified by centrifugation (20,000 g, 60 minutes, 4°C). SecA was purified from the soluble fraction as described for SecYEG but using TKM buffer with 30 (wash) or 300 mM (elution) imidazole. The eluate was further purified on a 25 ml bed volume column containing Q Sepharose fast flow resin equilibrated in TKM. After sample loading, a linear gradient of 50-1000 mM KCl was applied with SecA eluting at approximately 450 mM NaCl. SecA was subsequently purified with a HiLoad 26/600 Superdex 200 pg size exclusion column equilibrated in TKM, concentrated to approximately 500 µl with a 50 kDa centrifugal concentrator and stored at -80°C for future use.

GST-dark in pGEX-1 from our lab stock was transformed into BL21(DE3) cells and expressed as for SecYEG, using 1 mM IPTG for induction. Cells were resuspended in TK buffer (20 mM Tris-HCl pH 8.0 + 50 mM KCl), then cracked and centrifuged as for SecA. The supernatent was then loaded onto a GSTrap 4B column, washed with TK buffer, and eluted with 10 µM reduced glutathione in TK. Eluted protein was concentrated and stored at -80 ˚C.

11S in pBAD/*myc*-His C was transformed into BL21(DE3) and expressed in the same way as SecYEG, then resuspended in TKG (20 mM Tris pH 7.5, 50 mM KCl, 10 % glycerol) and cracked open and centrifuged as for SecA. Protein was purified by $Ni^{2+}$ affinity chromatography as above, using 50 mM imidazole for the wash step and 330 mM imidazole for the elution (in TKG). The imidazole was then exchanged away in a spin concentrator, and the pure protein concentrated and frozen at -80˚ C for later use.

#### ATPase assays

ATPase activity was followed by coupling ADP production to NADH oxidation using pyruvate kinase and lactate dehydrogenase (PK/LDH, Merck) in the presence of excess phophoenolpyruvate (PEP), then measuring reduction in NADH from its absorbance at 340 nm. Master mixes were assembled in TKM to give a final volume of 84 µl per well, with 1 µl PK/LDH, 5 µl PLs, and SecA, PEP and NADH to give final concentrations of 50 nM, 2 mM and 0.2 mM, respectively, when diluted to 100 µl. 84 µl master mix was added to clear, flat-bottomed 96-well plates (CoStar), and a baseline signal at 340 nm measured for 5 mins in a

BioTek Synergy Neo2 plate reader. Next, 1 µl 100 mM ATP was added manually, using a multichannel pipette, followed by 5 second double orbital shaking (559 cpm), and 20 min measurement of ATPase consumption in the absence of pre-protein. Finally, 15 µl pre-protein at 6.67 µM (to give 1 µM final) was added manually, followed by further shaking and measurement (stimulated activity). Slopes were determined by linear regression to the straight part of the line, then converted to ATP consumption by:

$$rate = \frac{-slope}{\varepsilon \times [SecA] \times l}$$

Where ε is the extinction coefficient for NADH at 340 nm (6,220 $M^{-1}.cm^{-1}$), [SecA] is the concentration of SecA (50 nM), and *l* is the path length (0.28 cm). Note that because the wells are cylindrical, the slightly higher [SecA] before pre-protein addition is precisely cancelled out by the shorter *l*. Eight simultaneous reactions were recorded per plate, and the identical repeats (4 per condition) averaged to produce a single value.

#### Transport assays

Transport assays were performed using the published Nanoluc system[1,2], wherein the large fragment of split NanoLuc (11S) is encapsulated within PLs with SecYEG in the membrane, and the small fragment (pep86) is incorporated into the pre-protein substrate. Upon transport of pre-protein into the vesicle, 11S and pep86 rapidly combine to form NanoLuc, which – when 11S is in excess and the substrate furimazine is provided – produces a luminescent signal proportional to the amount of pep86 imported.

NanoLuc measurements were performed on a BioTek Synergy Neo2 plate reader. Reaction master mixes were assembled at 25 ˚C to give a volume of 99 µl per well in TKM, such that when diluted to 125 µl each well contained: 0.625 µl furimazine, 0.1 % prionex, 0.1 $mg.ml^{-1}$ creatine kinase, 5 mM creatine phosphate, 20 µM GST-dark, 0.828 µl PLs and 500 µM SecA.

Immediately prior to measurement, 1 µl pre-protein at 250 µM was added to each measurement well in a white 96-well plate (Nunc), pre-warmed to 25 ˚C, followed by 99 µl master mix. After a 5 second double orbital shake (559 cpm), measurements were performed in kinetic mode (i.e. measuring absolute time from the start): 8 min luminescence measurement (to obtain background signal from unencapsulated 11S), injection of 25 µl ATP at 5 mM (in TKM), 5 second double orbital shake (559 cpm), and finally luminescence signal measurement (up to 50 minutes, but usually terminated early once all luminescence signals had plateaued).

Data processing was performed in pro Fit 7 (Quantumsoft), using the method established in Allen et al.[1]. First, the first 8 minutes were fitted to a single exponential + lag model, where $t_{app} = t$ - lag:

if ($t_{app}<0$) $$y = c$$

if ($t_{app}\geq0$) $$y = A(1 - e^{-t_{app} \times k}) + c$$

The resulting fit was extrapolated to cover the complete experiment time, tabulated, and subtracted from the data. Finally, the signal part of the corrected data was fitted to the same equation (with $t$=0 set to the injection time). Transport time is taken as lag from this fit, i.e. the minimum time for pep86 to reach the PL lumen.

#### Single-molecule data acquisition

Single-molecule Förster resonance energy transfer (smFRET) experiments were performed on a custom-built confocal epi-illuminated microscope[3] in a standard inverted-stage configuration. Molecules were excited in a pulsed interleaved excitation (PIE) regime using pulsed 560 nm (LDH-P-FA-560, Picoquant) and 640 nm (LDH-D-C-640, Picoquant) diode lasers which were alternatively pulsed at a combined frequency of 40 MHz by a Multichannel Picosecond Diode Laser Driver (Sepia PDL 828, Picoquant). The lasers were first combined using a dichroic mirror (86-394, Edmund Optics) before being coupled to an optical fiber

(PAF2A-11A, ThorLabs). Laser emission from the fiber was collimated (60FC-4-A11-01, Schäfter + Kirchhoff) before being reflected from a polychroic beam splitter (Di03-R405/488/561/635-t1-25x36m Semrock) into a 60x 1.2 NA water-immersion objective (UPLSAPO60XW, Olympus). Average powers for the donor and acceptor lasers at the objective were 75 µW and 35 µW respectively. Excitation light from the objective was focused 10 µm above the surface of the cover glass.

Light emitted from the sample was then collected by the same objective lens before being transmitted through the polychroic beam splitter. The emitted light was focused onto a 100 µM pinhole (P100D, ThorLabs) using an achromatic doublet lens with a focal length of 50 mm (AC254-050-A, ThorLabs). The pinhole spatially filtered photons outside of the desired observation volume. Light was then collimated using another achromatic lens of the same specifications before being seperated by wavelength into the donor and acceptor emission channels by a long-pass dichroic mirror (Di02-R635, Semrock). Bandpass filters removed residual excitation wavelength light in each channel (Donor: FF01-607/36-25, Semrock. Acceptor: FF01-670/30-25, Semrock) before it was focused by an achromatic doublet lens with a focal length of 50 mm (AC254-050-A, ThorLabs) onto single-photon avalanche diodes (SPCM-AQRH-14, Excelitas) with high timing resolution and low dark counts. Signals from the detectors are sent to a TCSPC data acquisition unit (Multiharp 150P, Picoquant).

#### Burst selection

We performed burst search and selection using the FRETBursts analysis software[4]. The background was assessed per each 60 s of acquisition, and bursts were identified as time periods during which the instantaneous photon count rate within a sliding window of 10 consecutive photons is at least 6 times higher than the background rate. Detected bursts which overlapped were fused. Bursts were selected if they include a minimum of 30 and a maximum of 1000 photons in total between all streams. The maximum photon threshold was included to filter out sparse aggregates of labelled SecYEG, which were significantly brighter than single-molecules.

#### mpH$^2$MM analysis

Bursts identified by FRETBursts were then converted into a format readable by the H2MM_C software[5]. All mpH$^2$MM calculations were performed within the Jupyter notebooks, available in supplementary dataset[6], using the Python package by Paul David Harris[7]. We used the mpH$^2$MM algorithm to test how well different state models describe the data.

#### Model selection

The modified Bayes Information Criterion (BIC')[5] was used for determining the number of states in the HMM model. The number of states for which BIC′ drops under 0.005 is generally considered reliable to describe the data with minimum parameters and this criterion was used throughout this study. Viterbi algorithm was used to find the most likely path between states based on the posterior probability. All data were best described by four states representing: i) dark donor, ii) dark acceptor, iii) low FRET (open state of SecYEG) and iv) high FRET (closed state of SecYEG) (see Main text Fig. 3).

#### Viterbi analysis

From the state path, photons were separated into dwells, each of which was assigned a duration, a mean $E_{raw}$, and a mean $S_{raw}$. This also allowed bursts to be classified by which and how many states they encompassed. As a measure of error, we used the weighted standard deviation and the weighted standard error of the $E_{raw}$ and $S_{raw}$ as a proxy for the standard error of the mpH$^2$MM model.

#### Dwell time error analysis

Analysis of the variability of subsets is a well-established method to assess the error of parameters. We repeated each experiment five times with freshly prepared proteoliposomes (see Supplementary Table 1, 2, 6 and 7) and calculated the average values together with a t-test to get the 90% confidence interval (90%CI). An example of the variability of the raw data is shown in Supplementary Fig. 16.

#### Burst variance analysis

As a qualitative test for FRET dynamics occurring within bursts, we used burst variance analysis (BVA)[8], which compares the expected variance in $E_{raw}$ based on shot noise (the static FRET semi-circle) against the actual variance in $E_{raw}$ in each individual burst. We performed BVA in the FRETbursts environment. To calculate the experimental burst wide standard deviation, we segmented each burst into consecutive (and nonoverlapping) windows of 5 photons each, and calculated the standard deviation of all windows within the burst. To further increase the statistical power of BVA, we segmented the $E_{raw}$ axis into bins, each centred on a given value of $E_{raw}$ and bearing an $E_{raw}$ bin width = 0.05.

#### Correction for the presence of apo SecYEG

During proteoliposome preparation, SecYEG complexes incorporate into the liposomes in a random fashion. This renders roughly 50% of molecules inaccessible[9], *i.e.* with the cytoplasmic loops facing inside the proteoliposome. These molecules cannot bind SecA and therefore remain in an ‘apo state’. To correct measured kinetics for the apo contribution, we simulated a mixture of transition rates of apo state with one unknown state over a physically reasonable range until we found a good agreement with measured data. The correction process pipeline is:

1. Mix apo and simulated transition rates:
    1. Simulate transition rates over a grid within a physically reasonable range.
    2. Mix simulated transition rates with apo in a 1:1 ratio.

2. Find the best agreement between simulated data and measured values to extract the corrected rates (i.e. transition rates before mixing with apo).

#### Simulation of transition rates

Stochastic conformational dynamics is often mathematically described using the "continuous-time Markov process", where the changes of state (conformation) of the system are called transitions. The probabilities associated with various state changes are called transition probabilities (P). The process is then characterised by a state space (S), a transition matrix (Q) describing the rates of particular transitions ($k$), and an initial state (or initial distribution) across the state space ($\pi_0$). Based on these parameters, we can determine a probability vector (π), which identifies the probability of the system being in a given state.

In our system, the state space consisted of two states, S={0, 1}. Here we used 0 to represent the open state of the LG and 1 for the closed state. We defined the rate of LG closing as $k_{01}$ as and the rate of LG opening as $k_{10}$.

$$Q = \begin{pmatrix} k_{00} & k_{01} \\ k_{10} & k_{11} \end{pmatrix} \quad \text{Eq.1}$$

$k_{00}$ and $k_{11}$ have no physical meaning, we can use them to justify a condition that a sum of each row of Q has to equal to zero, which means that $k_{00} = -k_{01}$ and $k_{11} = -k_{10}$. This condition is later needed to solve eq. 3. What we want to find is a probability vector π, which identifies the probability of the system being in a given state. In our example, π = (1, 0) would imply that the system stays all the time in state 0. On the other side, π = (0.5, 0.5) would imply an equal chance of being in both states. The relationship between Q and π is given by the following differential equation:

$$\frac{d\pi}{dt} = \pi Q \quad \text{Eq. 2}$$

What specifically is of interest for us is the steady state of a continuous Markov chain, in other words a state for which the derivative is 0:

$$\frac{d\pi}{dt} = \pi Q = 0 \qquad \text{Eq. 3}$$

We can quickly solve our differential equation analytically, using the matrix exponential:

$$\frac{d\pi}{dt} = \pi Q \Rightarrow \pi(t) = \pi(0)e^{Qt} \qquad \text{Eq. 4}$$

where the matrix exponential was computed using the python package scipy's linalg.expm function which implements an algorithm from Al-Mohy et al.[10]. Now, since we determined the probability vector π and we know the range of rates, we simulated all rates over a fine grid of opening and closing rates.

#### Mixing simulated rates with apo dynamics

In the next step, we mixed the simulated values with apo kinetics using weighted arithmetic average approach. Since opening and closing rates are independent, we mix the rates separately resulting in $k^{mix}_{\text{opening}}$ and $k^{mix}_{\text{closing}}$.

$$k^{mix}_{\text{opening}} = A^{APO}_{\text{opening}}\, k^{APO}_{\text{opening}} + A'_{\text{opening}}\, k'_{\text{opening}} \qquad \text{Eq. 5}$$

where

$$A^{APO}_{\text{opening}} = \frac{\pi^{APO}_{\text{opening}}}{\pi^{APO}_{\text{opening}} + \pi'_{\text{opening}}} \qquad \text{Eq. 6}$$

and

$$A'_{\text{opening}} = 1 - A^{APO}_{\text{opening}} \qquad \text{Eq. 7}$$

$\pi^{APO}_{\text{opening}}$ and $k^{APO}_{\text{opening}}$ were determined experimentally. $\pi'_{\text{opening}}$ was calculated for each simulated rate $k'_{\text{opening}}$ using the above-mentioned approach. Mixed closing rates were calculated the same way. Dwell times of each state are then defined as reciprocal values of a given rate. The Python script used for the generation of the correction matrix is attached under name "APO_correction.py".

#### Extracting the corrected rates

In the last step we recovered corrected rates by finding a minimum of the following expression:

$$\left(\left|k_{opening}^{measured} - k_{opening}^{mix}\right| + \left|k_{closing}^{measured} k_{closing}^{mix}\right|\right)/2$$

where $k_{opening}^{measured}$ *and* $k_{closing}^{measured}$ are the experimentally determined values. Calculated minima, showing the "apo-corrected" rates, for one repeat per technical repeat are depicted in Supplementary Fig. 5f-11f for SecYEG wild-type and Supplementary Fig. 13f-15f for SecYEG-PrlA4.

#### Calculation of simplified potential energy surface

The potential energy surface was estimated from transition state theory using the Kramers rate equation:

$$Q_{ij} = \begin{cases} \frac{D\sqrt{c_i c_{ij}}}{2\pi k_B T} e^{-\frac{G_{ij}^{\ddagger} - G_i}{k_B T}} & i \neq j \\ -\sum_{i \neq j} Q_{ij} & i = j \end{cases} \quad \text{Eq. 8}$$

where $Q_{ij}$ is the rate matrix, $D$ is the diffusion constant over an energy barrier; $c_i$ and $c_{ij}$ are the curvatures of the continuous free energy landscape at the energy minimum $G_i$ and the energy barrier $G^{\ddagger}_{ij}$ corresponding to the stable state *i* and the transition state from state *i* to state *j*, respectively. The minima and maxima of the energy landscape were interpreted as the free energy of the stable states and the transition states, respectively. Solving for the barrier heights we get:

$$G_{ij}^{\ddagger} = G_i - k_B T \log\left(2\pi Q_{ij} k_B T\right) - k_B T \log\left(D\sqrt{c_i c_{ij}}\right) \quad \text{Eq. 9}$$

We note that using eq. 9, we can only learn the exact energy of the barrier, $G^{\ddagger}_{ij}$, if we know $D$, $c_i$ and $c_{ij}$. However, $D$, $c_i$ and $c_{ij}$ are internal parameters of the system which are not easy to deduce. Moreover, the curvatures corresponding to different stable states are typically not equal. For example, proteins in folded states tend to be more rigid than in their unfolded states, leading to greater curvatures of the potential wells for the folded states than for the unfolded states. Nevertheless, in our system transitions occur between two well defined states and therefore we assume they do not significantly differ. We assume these parameters remain constant for all biochemical conditions in our experiments, therefore we can ignore the $D$, $c_i$ and $c_{ij}$.

#### Supplementary Figs.

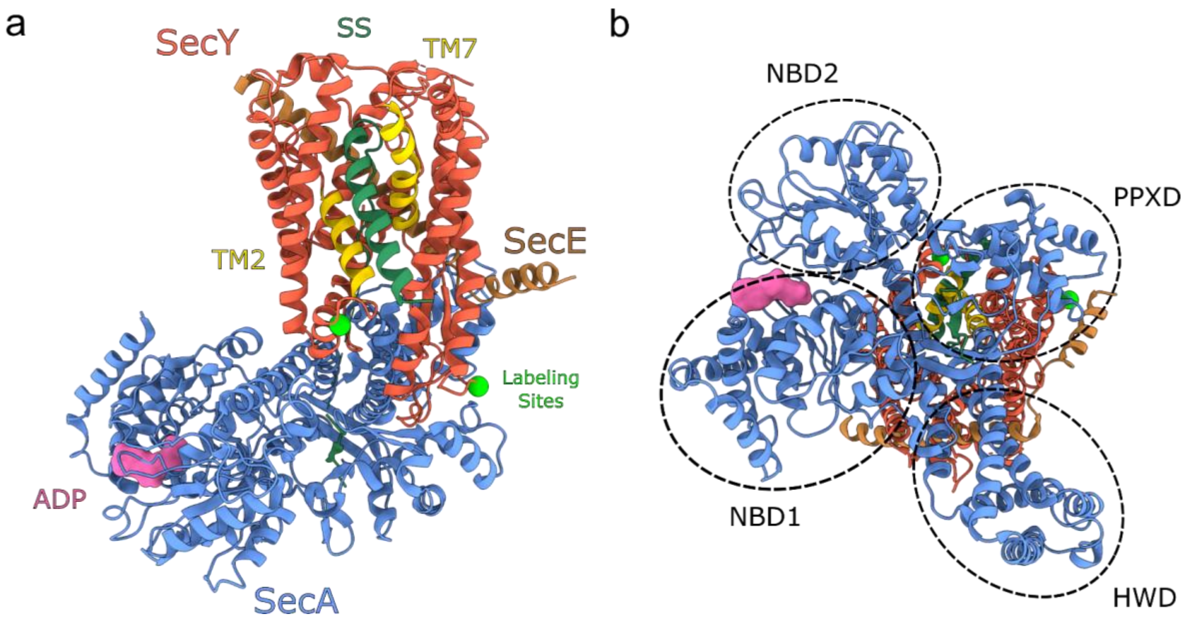


**Supplementary Fig. 1** - The structure of the SecYE:A complex from a) the side profile membrane view and b) the cytoplasmic view highlighting the nucleotide binding domains (NBD1+2), the pre-protein cross-linking domain (PPXD) and the helical wing domain (HWD). PDB: 6ITC[11] (structure determined in the absence of SecG).

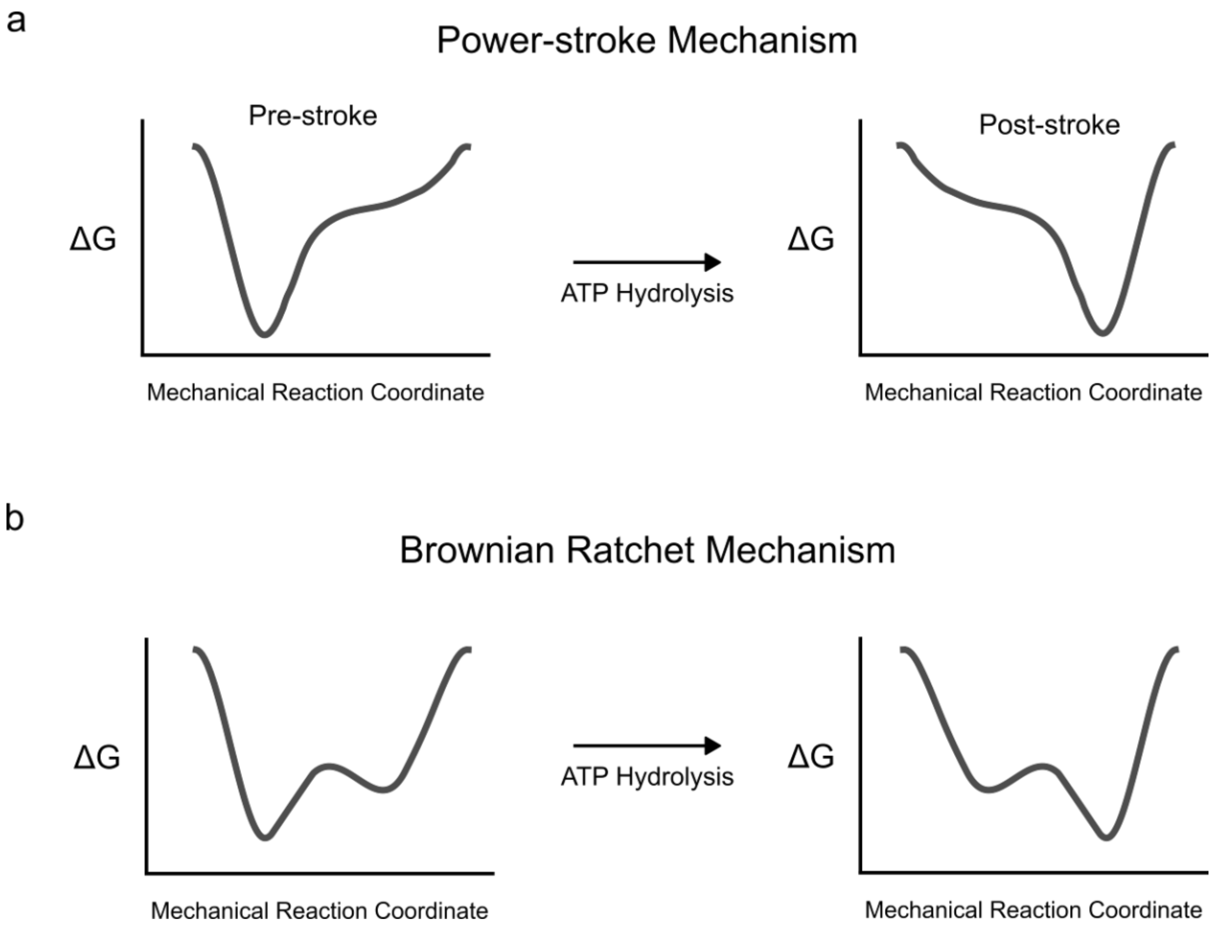


**Supplementary Fig. 2** - Mechanisms for protein translocation through the SecYEG:A complex illustrated by a free energy diagram across the mechanical reaction coordinate. a) Power-stroke mechanism: deep minima at either the pre-stroke or post-stroke position which directly correspond to the nucleotide state of the ATPase. b) Brownian ratchet mechanism: the nucleotide hydrolysis cycle biases a shallow energy profile towards certain conformations, which in turn undergo rapid interconversion between the available states due to thermal fluctuations.

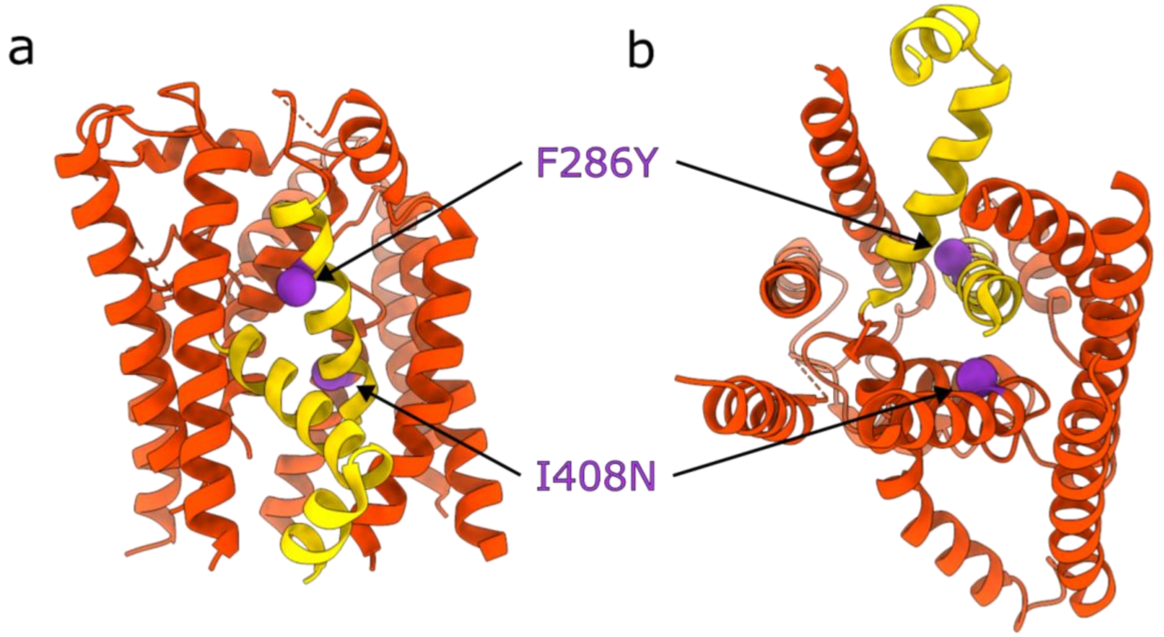


**Supplementary Fig. 3** - The structure of SecY showing the prlA4 mutations shown from a) the side profile membrane view and b) the cytoplasmic view. SecY is shown in red. The two prlA4 mutations F286Y and I408N are shown as purple space filled spheres. The two helices that comprise the lateral gate are shown in yellow (transmembrane helices 2 and 7). SecE, SecG and the cytoplasmic loops have been removed for clarity. PDB: 1RHZ[12].

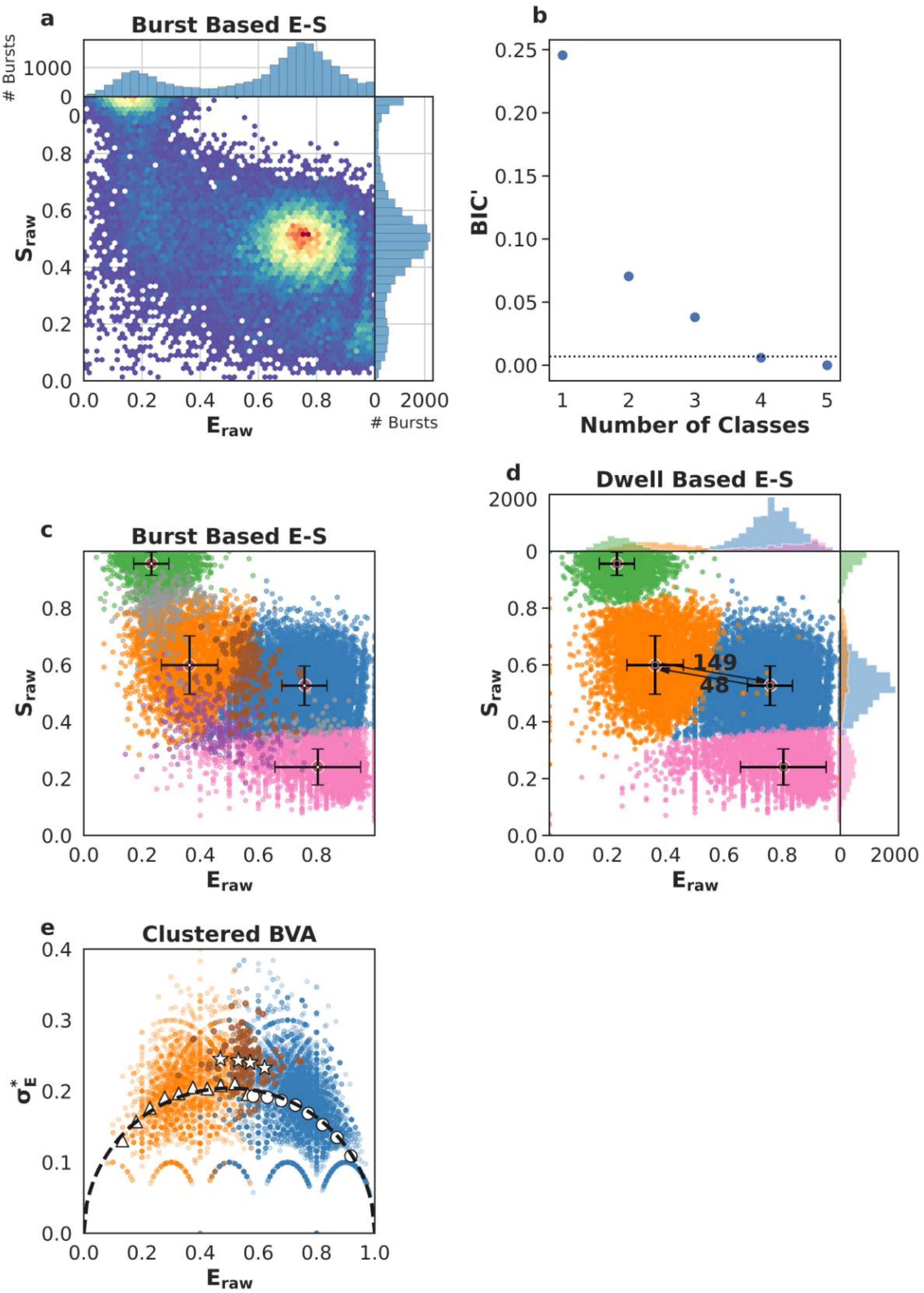
a
Burst Based E-S
### Bursts
$S_{raw}$
$E_{raw}$
b
BIC'
Number of Classes
c
Burst Based E-S
$S_{raw}$
$E_{raw}$
d
Dwell Based E-S
149
48
$S_{raw}$
$E_{raw}$
e
Clustered BVA
$\sigma_E^*$
$E_{raw}$

**Supplementary Fig. 4 –** Single-molecule data for one of five repeats of SecYEG apo (first column of data in Supplementary Table 1 and 2). a) Burst data plotting FRET efficiency ($E_{raw}$) against stoichiometry ($S_{raw}$). b) Modified Bayes Information Criterion (BIC') of different state-models indicated that four classes best describe the data (most likely state model BIC'<0.005). c) Using a burst based $E_{raw}$ $S_{raw}$ plot we interpreted the classes as the open state (orange, low $E_{raw}$), closed state (blue, high $E_{raw}$), dark acceptor (green, high $S_{raw}$), dark donor (pink, low $S_{raw}$), in this step, we also identified bursts representing transitions between conformational states (brown), transitions and photophysics (purple) and pure photophysics (grey). Crosses identify the average $E_{raw}$ $S_{raw}$ position for each state and the standard deviations. d) Dwell based $E_{raw}$ $S_{raw}$ plot showing the rate of interconversion ($s^{-1}$) between the open and closed states. e) Burst Variance Analysis (BVA) provided a qualitative confirmation of dynamic exchange between states and transition detection within bursts. Binned values representing bursts assigned to either the open (triangles) or the closed (circles) state, respectively, coincide with the static line (black), exhibiting no exchange on a millisecond time scale. On the other hand, bursts which mp$H_2$MM identified to capture dynamic interconversion (stars) were indeed found well above the static line, implying dynamics.

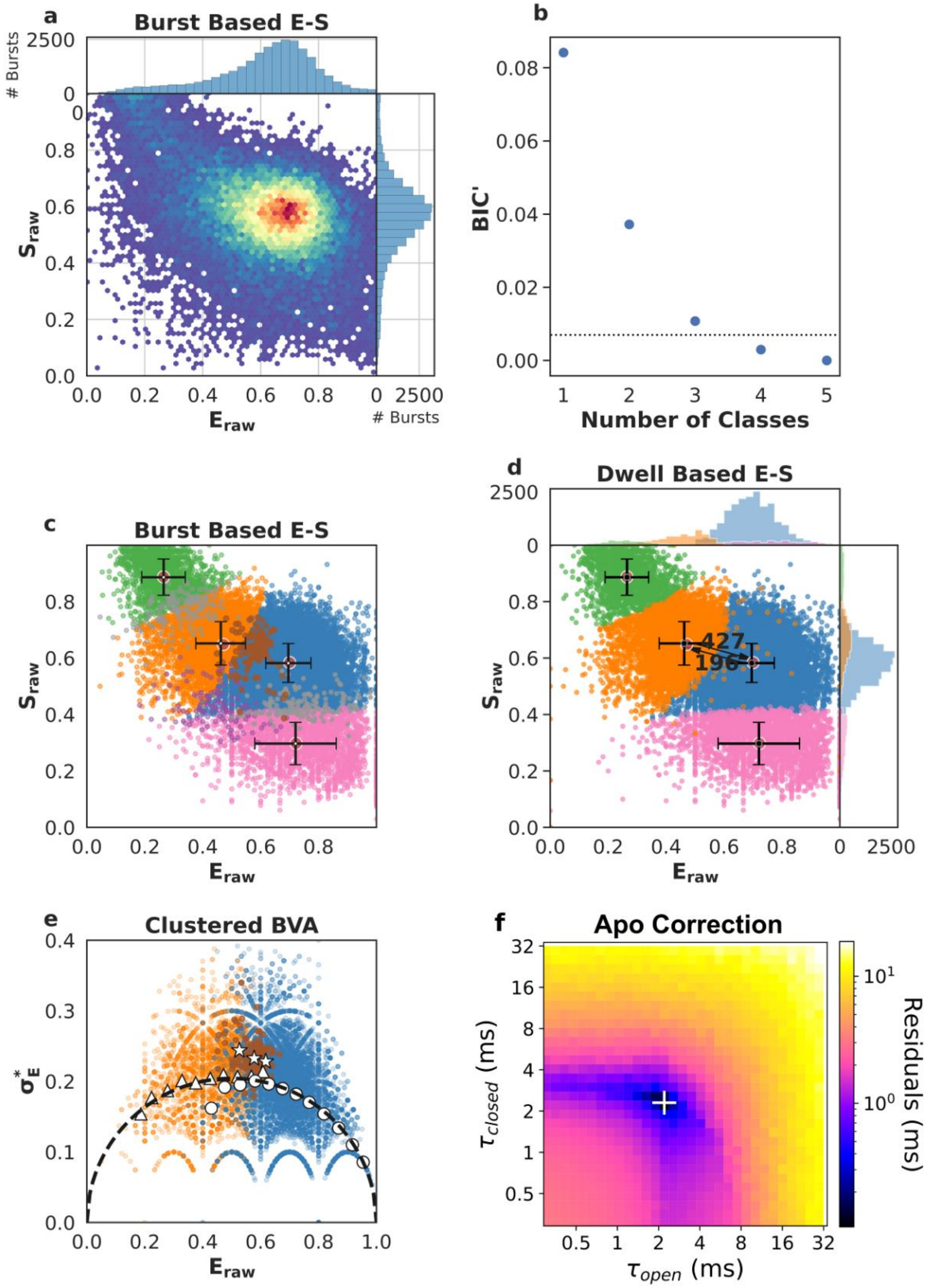

a
Burst Based E-S
### Bursts
$S_{raw}$
$E_{raw}$
### Bursts
b
BIC'
Number of Classes
c
Burst Based E-S
$S_{raw}$
$E_{raw}$
d
Dwell Based E-S
427
196
$S_{raw}$
$E_{raw}$
e
Clustered BVA
$\sigma_E^*$
$E_{raw}$
f
Apo Correction
$\tau_{closed}$ (ms)
$\tau_{open}$ (ms)
Residuals (ms)

**Supplementary Fig. 5** - Single-molecule data for one of five repeats of SecYEG + signal sequence (SecYEG+SS) (first column of data in Supplementary Table 1 and 2). a) Burst data plotting FRET efficiency ($E_{raw}$) against stoichiometry ($S_{raw}$). b) Modified Bayes Information Criterion (BIC’) of different state-models indicated that four classes best describe the data (most likely state model BIC’<0.005). c) Using a burst based $E_{raw}$ $S_{raw}$ plot we interpreted the classes as the open state (orange, low $E_{raw}$), closed state (blue, high $E_{raw}$), dark acceptor (green, high $S_{raw}$), dark donor (pink, low $S_{raw}$), in this step, we also identified bursts representing transitions between conformational states (brown), transitions and photophysics (purple) and pure photophysics (grey). Crosses identify the average $E_{raw}$ $S_{raw}$ position for each state and the standard deviations. d) Dwell based $E_{raw}$ $S_{raw}$ plot showing the rate of interconversion ($s^{-1}$) between the open and closed states. e) Burst Variance Analysis (BVA) provided a qualitative confirmation of dynamic exchange between states and transition detection within bursts. Binned values representing bursts assigned to either the open (triangles) or the closed (circles) state, respectively, coincide with the static line (black), exhibiting no exchange on a millisecond time scale. On the other hand, bursts which mpH$^2$MM identified to capture dynamic interconversion (stars) were indeed found well above the static line, implying dynamics. f) apo correction of extracted rates. The apo contribution is corrected for by finding the minimum difference between measured rates (panel d) and the numerical mixing of apo values with a range of opening and closing rates. The white '+' marks the recovered dwell times, i.e. the position of the lowest residuals.

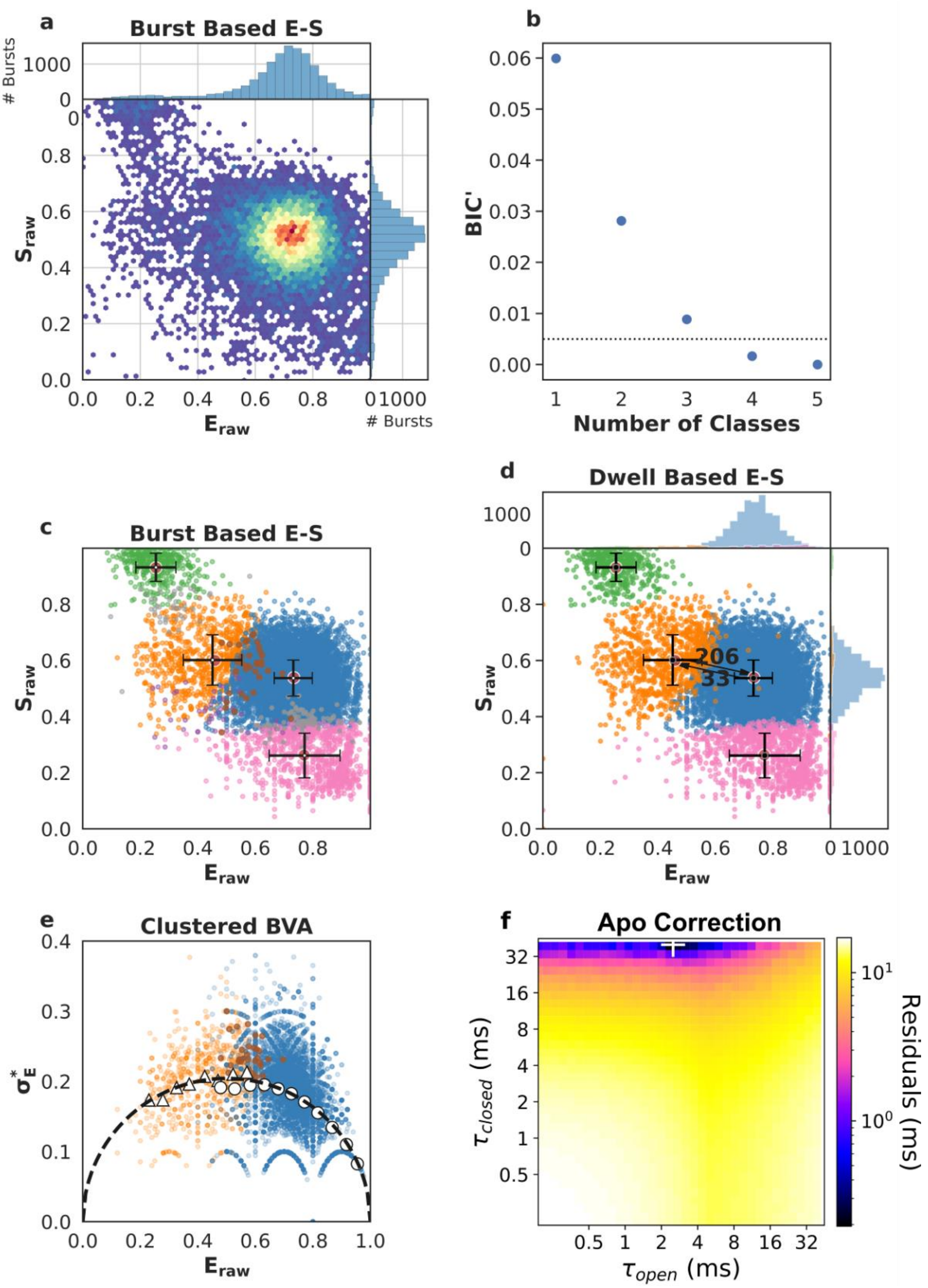
a
Burst Based E-S
### Bursts
$S_{raw}$
$E_{raw}$
b
BIC'
Number of Classes
c
Burst Based E-S
d
Dwell Based E-S
206
33
e
Clustered BVA
$\sigma_E^*$
f
Apo Correction
$\tau_{closed}$ (ms)
$\tau_{open}$ (ms)
Residuals (ms)

**Supplementary Fig. 6** - Single-molecule data for one of five repeats of SecYEG + SecA (SecYEG:A) (first column of data in Supplementary Table 1 and 2). a) Burst data plotting FRET efficiency ($E_{raw}$) against stoichiometry ($S_{raw}$). b) Modified Bayes Information Criterion (BIC') of different state-models indicated that four classes best describe the data (most likely state model BIC'<0.005). c) Using a burst based $E_{raw}$ $S_{raw}$ plot we interpreted the classes as the open state (orange, low $E_{raw}$), closed state (blue, high $E_{raw}$), dark acceptor (green, high $S_{raw}$), dark donor (pink, low $S_{raw}$), in this step, we also identified bursts representing transitions between conformational states (brown), transitions and photophysics (purple) and pure photophysics (grey). Crosses identify the average $E_{raw}$ $S_{raw}$ position for each state and the standard deviations. d) Dwell based $E_{raw}$ $S_{raw}$ plot showing the rate of interconversion ($s^{-1}$) between the open and closed states. e) Burst Variance Analysis (BVA) provided a qualitative confirmation of dynamic exchange between states and transition detection within bursts. Binned values representing bursts assigned to either the open (triangles) or the closed (circles) state, respectively, coincide with the static line (black), exhibiting no exchange on a millisecond time scale. On the other hand, bursts which mp$H_2$MM identified to capture dynamic interconversion (stars) were indeed found well above the static line, implying dynamics. f) apo correction of extracted rates. The apo contribution is corrected for by finding the minimum difference between measured rates (panel d) and the numerical mixing of apo values with a range of opening and closing rates. The white '+' marks the recovered dwell times, i.e. the position of the lowest residuals.

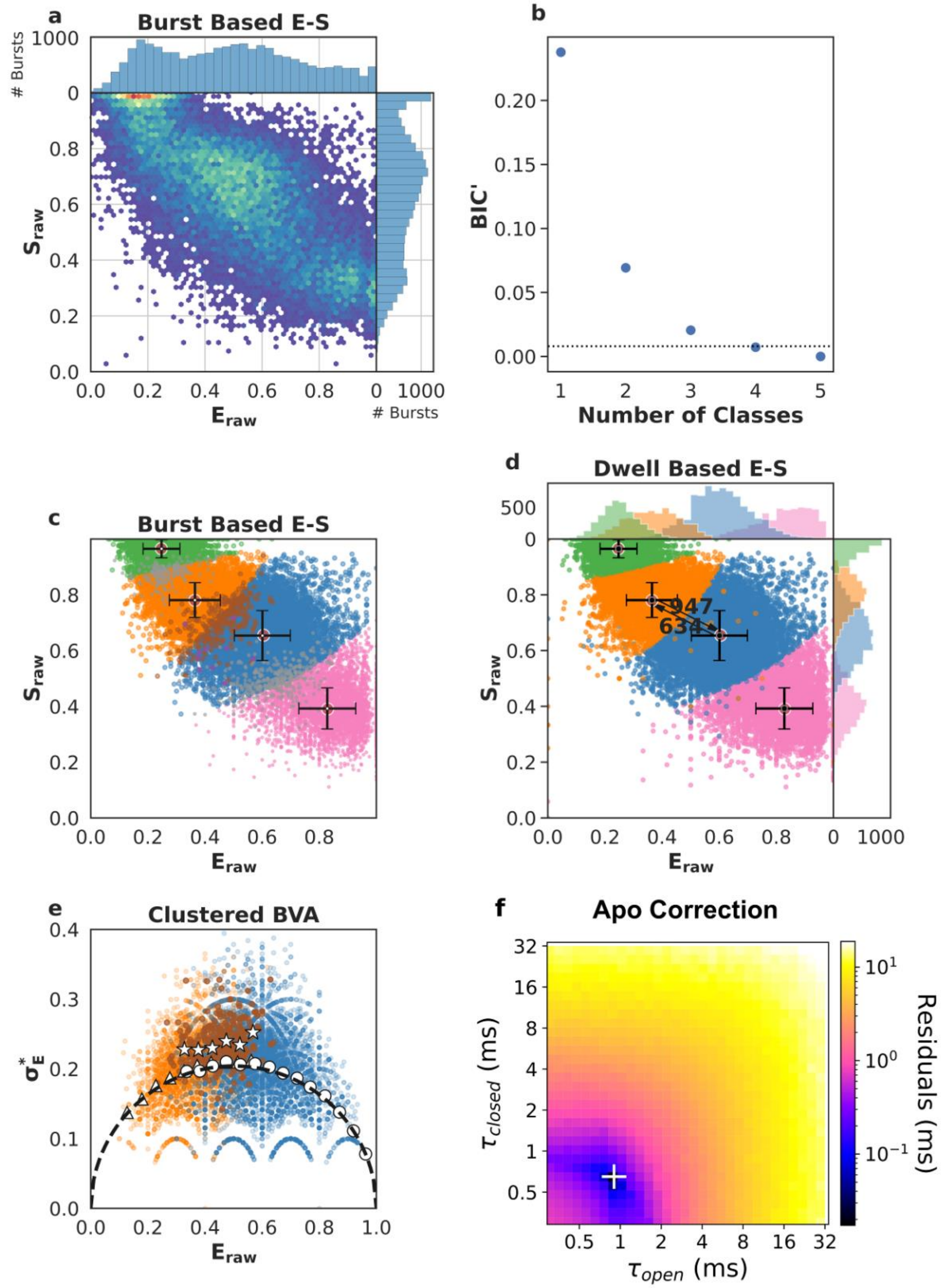
a
Burst Based E-S
### Bursts
1000
0
0.8
0.6
0.4
0.2
0.0
$S_{raw}$
0.0 0.2 0.4 0.6 0.8
$E_{raw}$
0 1000
### Bursts
b
0.20
0.15
0.10
0.05
0.00
BIC'
1 2 3 4 5
Number of Classes
c
Burst Based E-S
$S_{raw}$
$E_{raw}$
d
Dwell Based E-S
500
947
634
$S_{raw}$
$E_{raw}$
e
Clustered BVA
$\sigma_E^*$
$E_{raw}$
f
Apo Correction
$\tau_{closed}$ (ms)
$\tau_{open}$ (ms)
Residuals (ms)
$10^1$
$10^0$
$10^{-1}$

**Supplementary Fig. 7** - Single-molecule data for one of five repeats of SecYEG + SecA + ATPγS (SecYEG:A:ATPγS) (first column of data in Supplementary Table 1 and 2). a) Burst data plotting FRET efficiency ($E_{raw}$) against stoichiometry ($S_{raw}$). b) Modified Bayes Information Criterion (BIC') of different state-models indicated that four classes best describe the data (most likely state model BIC'<0.005). c) Using a burst based $E_{raw}$ $S_{raw}$ plot we interpreted the classes as the open state (orange, low $E_{raw}$), closed state (blue, high $E_{raw}$), dark acceptor (green, high $S_{raw}$), dark donor (pink, low $S_{raw}$), in this step, we also identified bursts representing transitions between conformational states (brown), transitions and photophysics (purple) and pure photophysics (grey). Crosses identify the average $E_{raw}$ $S_{raw}$ position for each state and the standard deviations. d) Dwell based $E_{raw}$ $S_{raw}$ plot showing the rate of interconversion ($s^{-1}$) between the open and closed states. e) Burst Variance Analysis (BVA) provided a qualitative confirmation of dynamic exchange between states and transition detection within bursts. Binned values representing bursts assigned to either the open (triangles) or the closed (circles) state, respectively, coincide with the static line (black), exhibiting no exchange on a millisecond time scale. On the other hand, bursts which mp$H_2$MM identified to capture dynamic interconversion (stars) were indeed found well above the static line, implying dynamics. f) apo correction of extracted rates. The apo contribution is corrected for by finding the minimum difference between measured rates (panel d) and the numerical mixing of apo values with a range of opening and closing rates. The white '+' marks the recovered dwell times, i.e. the position of the lowest residuals.

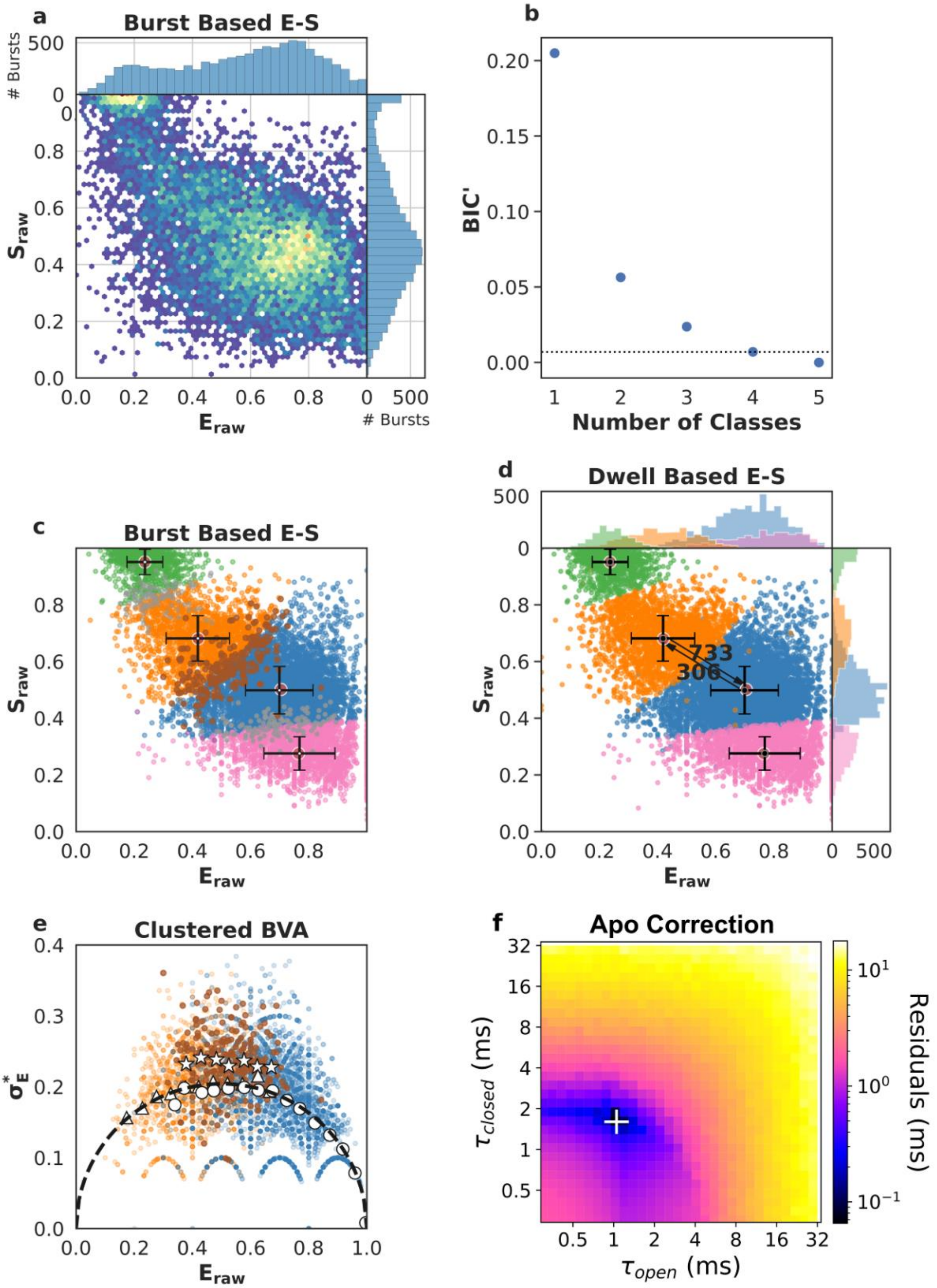
a
Burst Based E-S
### Bursts
$S_{raw}$
$E_{raw}$
### Bursts
b
BIC'
Number of Classes
c
Burst Based E-S
$S_{raw}$
$E_{raw}$
d
Dwell Based E-S
733
306
$S_{raw}$
$E_{raw}$
e
Clustered BVA
$\sigma_E^*$
$E_{raw}$
f
Apo Correction
$\tau_{closed}$ (ms)
$\tau_{open}$ (ms)
Residuals (ms)

**Supplementary Fig. 8** - Single-molecule data for one of five repeats of SecYEG + SecA + ADP·$AlF_x$ (SecYEG:A:ADP·$AlF_x$) (first column of data in Supplementary Table 1 and 2). a) Burst data plotting FRET efficiency ($E_{raw}$) against stoichiometry ($S_{raw}$). b) Modified Bayes Information Criterion (BIC') of different state-models indicated that four classes best describe the data (most likely state model BIC'<0.005). c) Using a burst based $E_{raw}$ $S_{raw}$ plot we interpreted the classes as the open state (orange, low $E_{raw}$), closed state (blue, high $E_{raw}$), dark acceptor (green, high $S_{raw}$), dark donor (pink, low $S_{raw}$), in this step, we also identified bursts representing transitions between conformational states (brown), transitions and photophysics (purple) and pure photophysics (grey). Crosses identify the average $E_{raw}$ $S_{raw}$ position for each state and the standard deviations. d) Dwell based $E_{raw}$ $S_{raw}$ plot showing the rate of interconversion ($s^{-1}$) between the open and closed states. e) Burst Variance Analysis (BVA) provided a qualitative confirmation of dynamic exchange between states and transition detection within bursts. Binned values representing bursts assigned to either the open (triangles) or the closed (circles) state, respectively, coincide with the static line (black), exhibiting no exchange on a millisecond time scale. On the other hand, bursts which mp$H_2$MM identified to capture dynamic interconversion (stars) were indeed found well above the static line, implying dynamics. f) apo correction of extracted rates. The apo contribution is corrected for by finding the minimum difference between measured rates (panel d) and the numerical mixing of apo values with a range of opening and closing rates. The white '+' marks the recovered dwell times, i.e. the position of the lowest residuals.

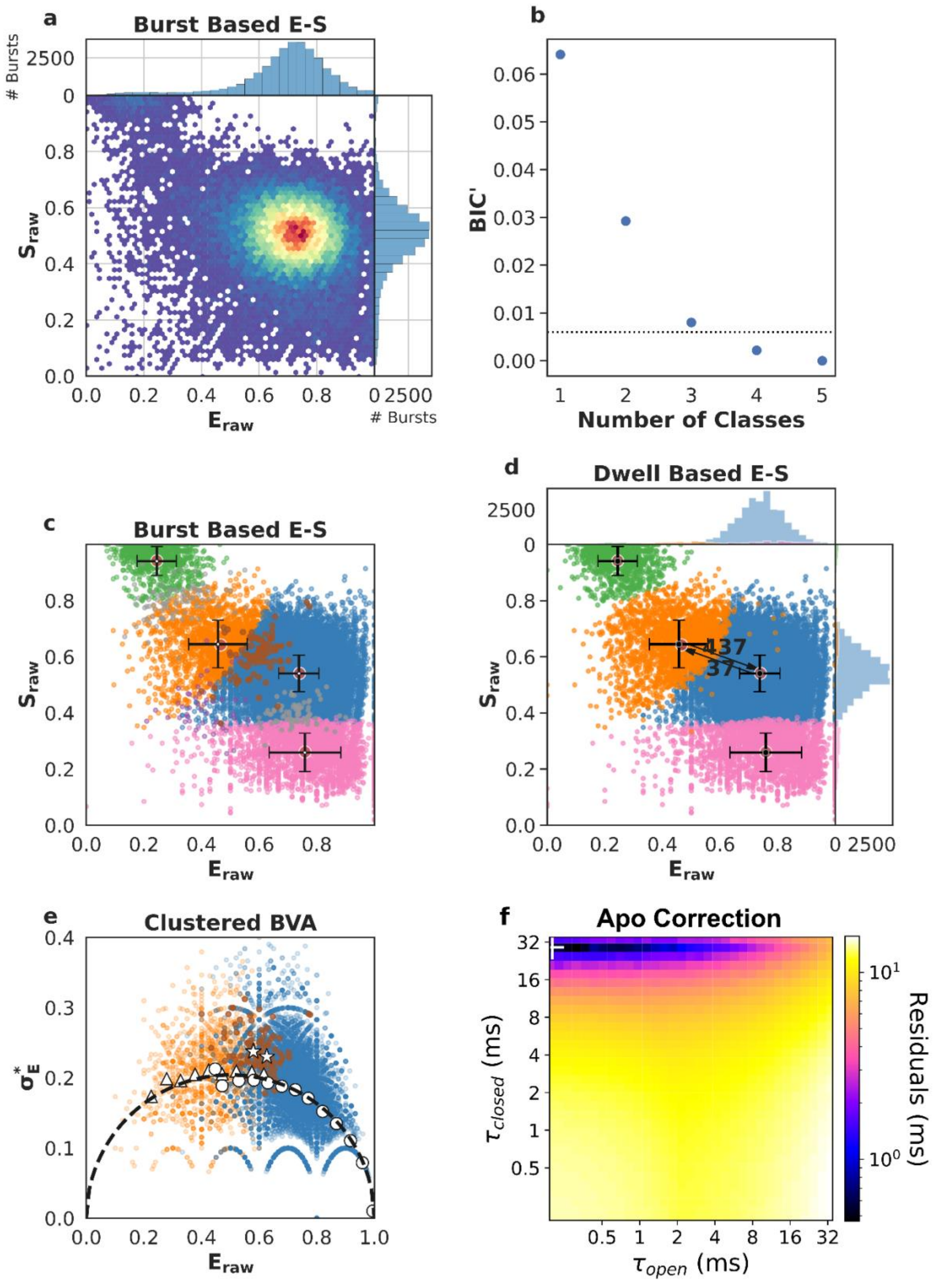
a
Burst Based E-S
### Bursts
2500
0
0.8
0.6
0.4
0.2
0.0
$S_{raw}$
0.0 0.2 0.4 0.6 0.8
$E_{raw}$
0 2500
### Bursts
b
BIC'
0.06
0.05
0.04
0.03
0.02
0.01
0.00
1 2 3 4 5
Number of Classes
c
Burst Based E-S
d
Dwell Based E-S
437
37
e
Clustered BVA
$\sigma_E^*$
0.4
0.3
0.2
0.1
0.0
0.0 0.2 0.4 0.6 0.8 1.0
f
Apo Correction
$\tau_{closed}$ (ms)
32
16
8
4
2
1
0.5
0.5 1 2 4 8 16 32
$\tau_{open}$ (ms)
$10^1$
$10^0$
Residuals (ms)

**Supplementary Fig. 9** - Single-molecule data for one of five repeats of SecYEG + SecA + ADP (SecYEG:A:ADP) (first column of data in Supplementary Table 1 and 2). a) Burst data plotting FRET efficiency ($E_{raw}$) against stoichiometry ($S_{raw}$). b) Modified Bayes Information Criterion (BIC') of different state-models indicated that four classes best describe the data (most likely state model BIC'<0.005). c) Using a burst based $E_{raw}$ $S_{raw}$ plot we interpreted the classes as the open state (orange, low $E_{raw}$), closed state (blue, high $E_{raw}$), dark acceptor (green, high $S_{raw}$), dark donor (pink, low $S_{raw}$), in this step, we also identified bursts representing transitions between conformational states (brown), transitions and photophysics (purple) and pure photophysics (grey). Crosses identify the average $E_{raw}$ $S_{raw}$ position for each state and the standard deviations. d) Dwell based $E_{raw}$ $S_{raw}$ plot showing the rate of interconversion ($s^{-1}$) between the open and closed states. e) Burst Variance Analysis (BVA) provided a qualitative confirmation of dynamic exchange between states and transition detection within bursts. Binned values representing bursts assigned to either the open (triangles) or the closed (circles) state, respectively, coincide with the static line (black), exhibiting no exchange on a millisecond time scale. On the other hand, bursts which mp$H_2$MM identified to capture dynamic interconversion (stars) were indeed found well above the static line, implying dynamics. f) apo correction of extracted rates. The apo contribution is corrected for by finding the minimum difference between measured rates (panel d) and the numerical mixing of apo values with a range of opening and closing rates. The white '+' marks the recovered dwell times, i.e. the position of the lowest residuals.

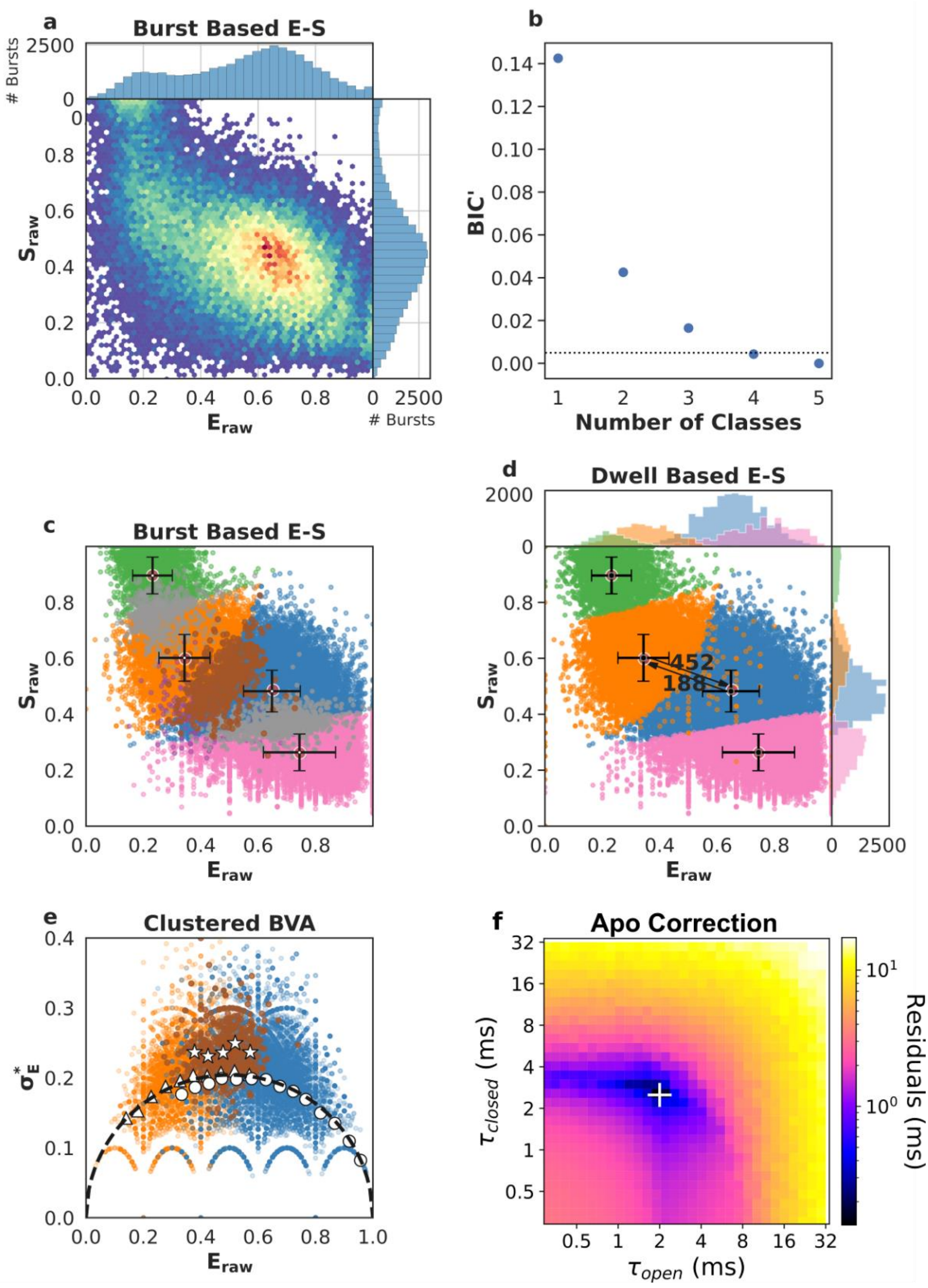
a
Burst Based E-S
### Bursts
$S_{raw}$
$E_{raw}$
b
BIC'
Number of Classes
c
Burst Based E-S
d
Dwell Based E-S
452
188
e
Clustered BVA
$\sigma_E^*$
f
Apo Correction
$\tau_{closed}$ (ms)
$\tau_{open}$ (ms)
Residuals (ms)

**Supplementary Fig. 10** - Single-molecule data for one of five repeats of SecYEG + SecA + ATP (SecYEG:A:ATP) (first column of data in Supplementary Table 1 and 2). a) Burst data plotting FRET efficiency ($E_{raw}$) against stoichiometry ($S_{raw}$). b) Modified Bayes Information Criterion (BIC') of different state-models indicated that four classes best describe the data (most likely state model BIC'<0.005). c) Using a burst based $E_{raw}$ $S_{raw}$ plot we interpreted the classes as the open state (orange, low $E_{raw}$), closed state (blue, high $E_{raw}$), dark acceptor (green, high $S_{raw}$), dark donor (pink, low $S_{raw}$), in this step, we also identified bursts representing transitions between conformational states (brown), transitions and photophysics (purple) and pure photophysics (grey). Crosses identify the average $E_{raw}$ $S_{raw}$ position for each state and the standard deviations. d) Dwell based $E_{raw}$ $S_{raw}$ plot showing the rate of interconversion ($s^{-1}$) between the open and closed states. e) Burst Variance Analysis (BVA) provided a qualitative confirmation of dynamic exchange between states and transition detection within bursts. Binned values representing bursts assigned to either the open (triangles) or the closed (circles) state, respectively, coincide with the static line (black), exhibiting no exchange on a millisecond time scale. On the other hand, bursts which mp$H_2$MM identified to capture dynamic interconversion (stars) were indeed found well above the static line, implying dynamics. f) apo correction of extracted rates. The apo contribution is corrected for by finding the minimum difference between measured rates (panel d) and the numerical mixing of apo values with a range of opening and closing rates. The white '+' marks the recovered dwell times, i.e. the position of the lowest residuals.

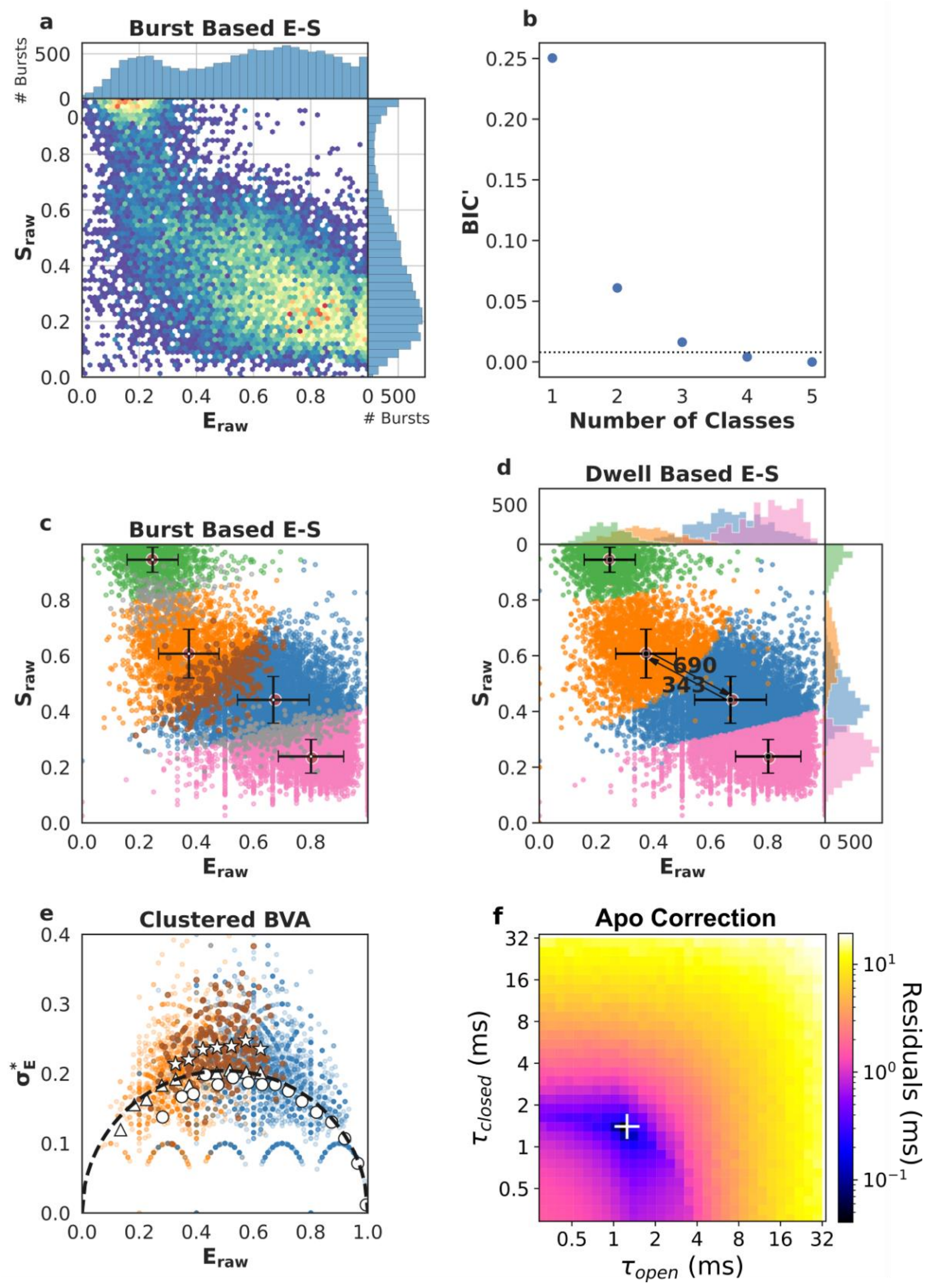

a
Burst Based E-S
### Bursts
Sraw
Eraw
b
BIC'
Number of Classes
c
Burst Based E-S
d
Dwell Based E-S
690
343
e
Clustered BVA
σ*E
f
Apo Correction
τclosed (ms)
τopen (ms)
Residuals (ms)

**Supplementary Fig. 11** - Single-molecule data for one of five repeats of SecYEG + SecA + ATP + proSpy (SecYEG:A:proSpy:ATP) (first column of data in Supplementary Table 1 and 2). a) Burst data plotting FRET efficiency ($E_{raw}$) against stoichiometry ($S_{raw}$). b) Modified Bayes Information Criterion (BIC') of different state-models indicated that four classes best describe the data (most likely state model BIC'<0.005). c) Using a burst based $E_{raw}$ $S_{raw}$ plot we interpreted the classes as the open state (orange, low $E_{raw}$), closed state (blue, high $E_{raw}$), dark acceptor (green, high $S_{raw}$), dark donor (pink, low $S_{raw}$), in this step, we also identified bursts representing transitions between conformational states (brown), transitions and photophysics (purple) and pure photophysics (grey). Crosses identify the average $E_{raw}$ $S_{raw}$ position for each state and the standard deviations. d) Dwell based $E_{raw}$ $S_{raw}$ plot showing the rate of interconversion ($s^{-1}$) between the open and closed states. e) Burst Variance Analysis (BVA) provided a qualitative confirmation of dynamic exchange between states and transition detection within bursts. Binned values representing bursts assigned to either the open (triangles) or the closed (circles) state, respectively, coincide with the static line (black), exhibiting no exchange on a millisecond time scale. On the other hand, bursts which mp$H_2$MM identified to capture dynamic interconversion (stars) were indeed found well above the static line, implying dynamics. f) apo correction of extracted rates. The apo contribution is corrected for by finding the minimum difference between measured rates (panel d) and the numerical mixing of apo values with a range of opening and closing rates. The white '+' marks the recovered dwell times, i.e. the position of the lowest residuals.

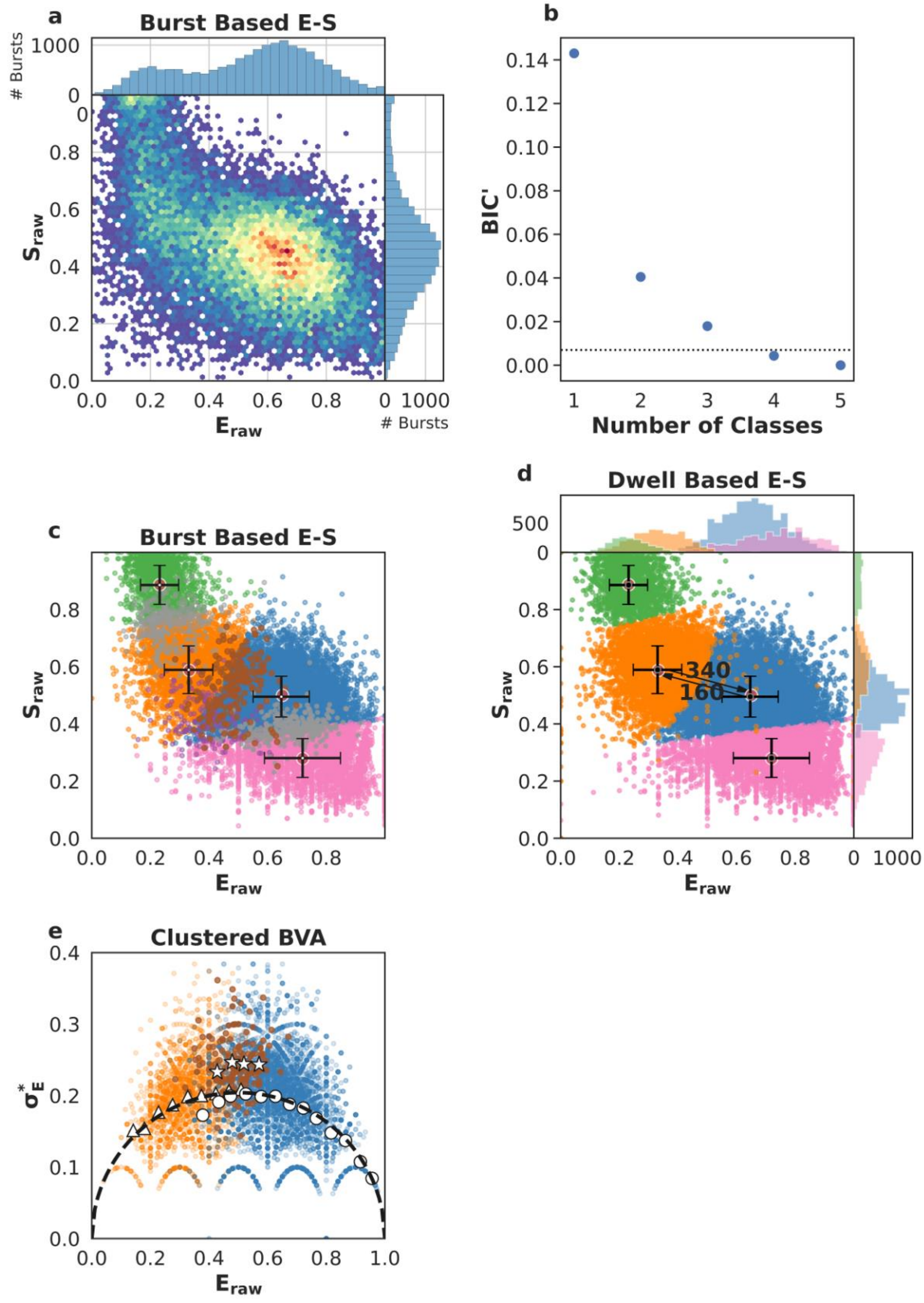
a
Burst Based E-S
### Bursts
1000
0
$S_{raw}$
$E_{raw}$
### Bursts
b
BIC'
Number of Classes
c
Burst Based E-S
$S_{raw}$
$E_{raw}$
d
Dwell Based E-S
500
340
160
$S_{raw}$
$E_{raw}$
e
Clustered BVA
$\sigma_E^*$
$E_{raw}$

**Supplementary Fig. 12** - Single-molecule data for one of five repeats of SecYEG-PrlA4 apo (first column of data in Supplementary Table 6 and 7). a) Burst data plotting FRET efficiency ($E_{raw}$) against stoichiometry ($S_{raw}$). b) Modified Bayes Information Criterion (BIC') of different state-models indicated that four classes best describe the data (most likely state model BIC'<0.005). c) Using a burst based $E_{raw}$ $S_{raw}$ plot we interpreted the classes as the open state (orange, low $E_{raw}$), closed state (blue, high $E_{raw}$), dark acceptor (green, high $S_{raw}$), dark donor (pink, low $S_{raw}$), in this step, we also identified bursts representing transitions between conformational states (brown), transitions and photophysics (purple) and pure photophysics (grey). Crosses identify the average $E_{raw}$ $S_{raw}$ position for each state and the standard deviations. d) Dwell based $E_{raw}$ $S_{raw}$ plot showing the rate of interconversion ($s^{-1}$) between the open and closed states. e) Burst Variance Analysis (BVA) provided a qualitative confirmation of dynamic exchange between states and transition detection within bursts. Binned values representing bursts assigned to either the open (triangles) or the closed (circles) state, respectively, coincide with the static line (black), exhibiting no exchange on a millisecond time scale. On the other hand, bursts which mp$H_2$MM identified to capture dynamic interconversion (stars) were indeed found well above the static line, implying dynamics.

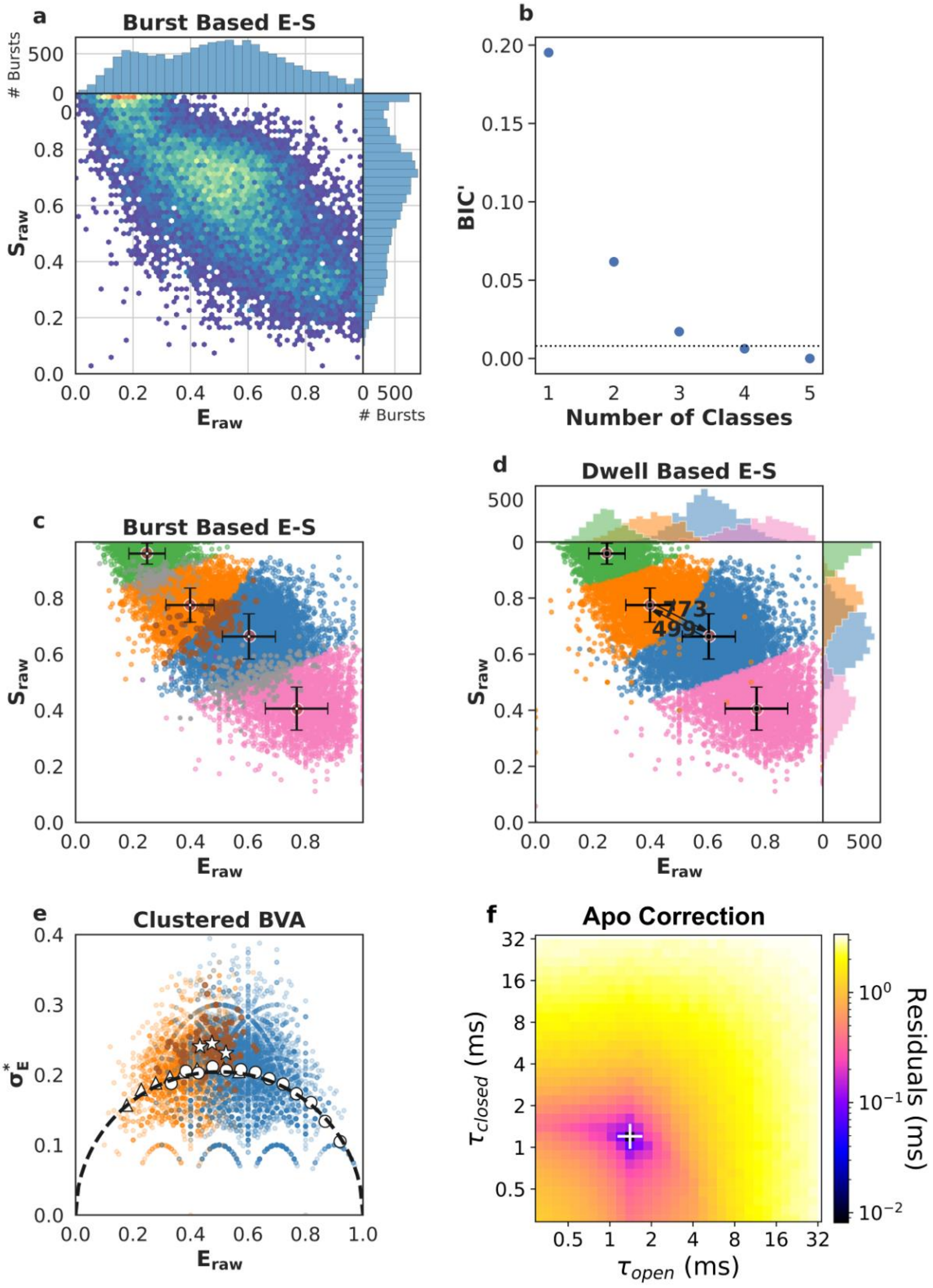
a
Burst Based E-S
### Bursts
500
0
$S_{raw}$
0.0
0.2
0.4
0.6
0.8
$E_{raw}$
0 500
### Bursts
b
BIC'
0.00
0.05
0.10
0.15
0.20
1 2 3 4 5
Number of Classes
c
Burst Based E-S
$S_{raw}$
$E_{raw}$
d
Dwell Based E-S
500
773
499
$S_{raw}$
$E_{raw}$
0 500
e
Clustered BVA
$\sigma_E^*$
0.0
0.1
0.2
0.3
0.4
0.0 0.2 0.4 0.6 0.8 1.0
$E_{raw}$
f
Apo Correction
$\tau_{closed}$ (ms)
0.5 1 2 4 8 16 32
$\tau_{open}$ (ms)
$10^0$
$10^{-1}$
$10^{-2}$
Residuals (ms)

**Supplementary Fig. 13** - Single-molecule data for one of five repeats of SecYEG-PrlA4 + signal sequence (SecYEG-PrlA4:SS) (first column of data in Supplementary Table 6 and 7). a) Burst data plotting FRET efficiency ($E_{raw}$) against stoichiometry ($S_{raw}$). b) Modified Bayes Information Criterion (BIC') of different state-models indicated that four classes best describe the data (most likely state model BIC'<0.005). c) Using a burst based $E_{raw}$ $S_{raw}$ plot we interpreted the classes as the open state (orange, low $E_{raw}$), closed state (blue, high $E_{raw}$), dark acceptor (green, high $S_{raw}$), dark donor (pink, low $S_{raw}$), in this step, we also identified bursts representing transitions between conformational states (brown), transitions and photophysics (purple) and pure photophysics (grey). Crosses identify the average $E_{raw}$ $S_{raw}$ position for each state and the standard deviations. d) Dwell based $E_{raw}$ $S_{raw}$ plot showing the rate of interconversion ($s^{-1}$) between the open and closed states. e) Burst Variance Analysis (BVA) provided a qualitative confirmation of dynamic exchange between states and transition detection within bursts. Binned values representing bursts assigned to either the open (triangles) or the closed (circles) state, respectively, coincide with the static line (black), exhibiting no exchange on a millisecond time scale. On the other hand, bursts which mp$H_2$MM identified to capture dynamic interconversion (stars) were indeed found well above the static line, implying dynamics. f) apo correction of extracted rates. The apo contribution is corrected for by finding the minimum difference between measured rates (panel d) and the numerical mixing of apo values with a range of opening and closing rates. The white '+' marks the recovered dwell times, i.e. the position of the lowest residuals.

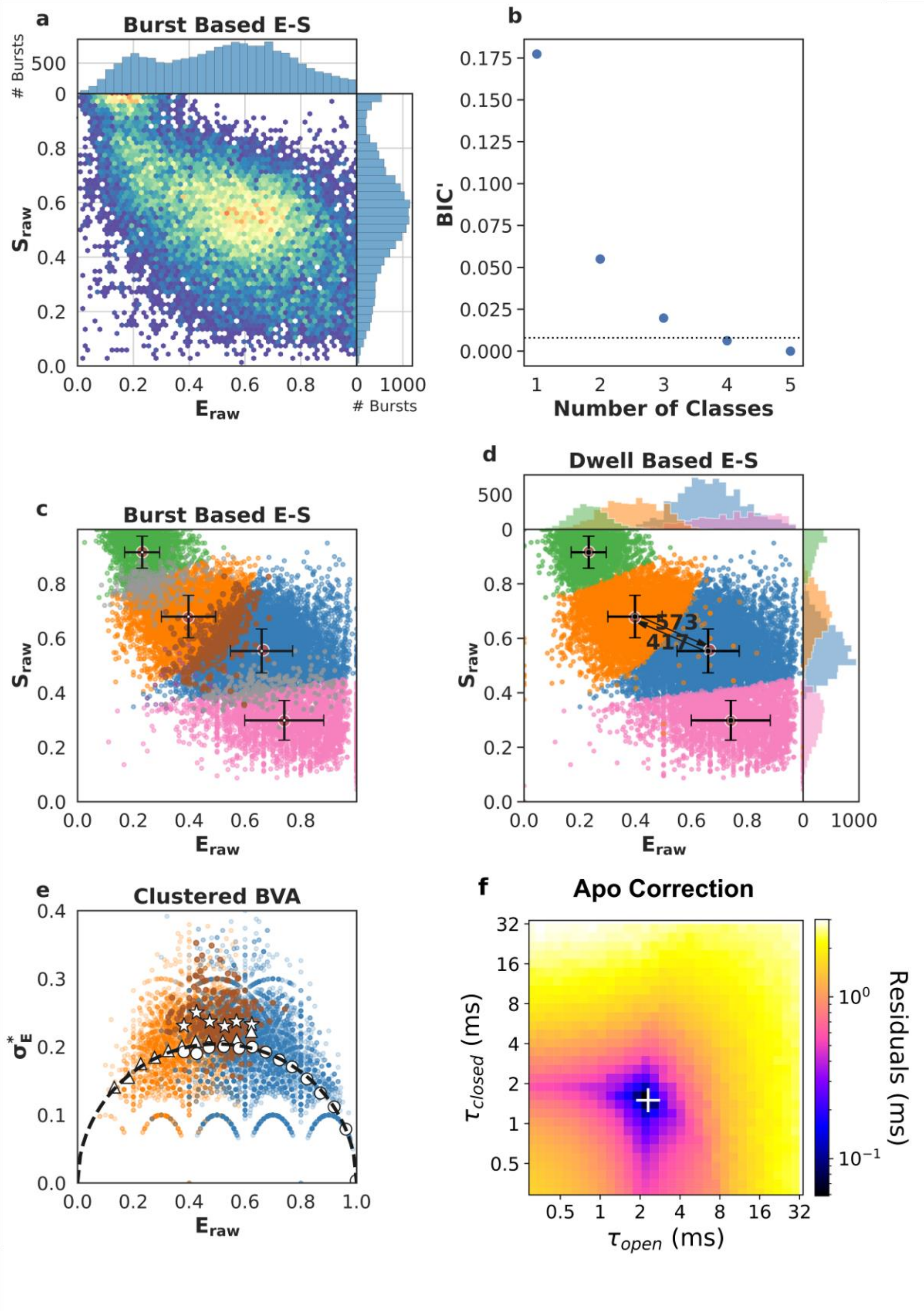
a
Burst Based E-S
### Bursts
500
0
0.8
0.6
0.4
0.2
0.0
S_raw
0.0 0.2 0.4 0.6 0.8
E_raw
0 1000
### Bursts
b
0.175
0.150
0.125
0.100
0.075
0.050
0.025
0.000
BIC'
1 2 3 4 5
Number of Classes
c
Burst Based E-S
S_raw
E_raw
d
Dwell Based E-S
500
0
573
417
S_raw
E_raw
0 1000
e
Clustered BVA
0.4
0.3
0.2
0.1
0.0
σ*_E
0.0 0.2 0.4 0.6 0.8 1.0
E_raw
f
Apo Correction
32
16
8
4
2
1
0.5
τ_closed (ms)
0.5 1 2 4 8 16 32
τ_open (ms)
10^0
10^-1
Residuals (ms)

**Supplementary Fig. 14** - Single-molecule data for one of five repeats of SecYEG-PrlA4 + SecA (SecYEG-PrlA4:A) (first column of data in Supplementary Table 6 and 7). a) Burst data plotting FRET efficiency ($E_{raw}$) against stoichiometry ($S_{raw}$). b) Modified Bayes Information Criterion (BIC') of different state-models indicated that four classes best describe the data (most likely state model BIC'<0.005). c) Using a burst based $E_{raw}$ $S_{raw}$ plot we interpreted the classes as the open state (orange, low $E_{raw}$), closed state (blue, high $E_{raw}$), dark acceptor (green, high $S_{raw}$), dark donor (pink, low $S_{raw}$), in this step, we also identified bursts representing transitions between conformational states (brown), transitions and photophysics (purple) and pure photophysics (grey). Crosses identify the average $E_{raw}$ $S_{raw}$ position for each state and the standard deviations. d) Dwell based $E_{raw}$ $S_{raw}$ plot showing the rate of interconversion ($s^{-1}$) between the open and closed states. e) Burst Variance Analysis (BVA) provided a qualitative confirmation of dynamic exchange between states and transition detection within bursts. Binned values representing bursts assigned to either the open (triangles) or the closed (circles) state, respectively, coincide with the static line (black), exhibiting no exchange on a millisecond time scale. On the other hand, bursts which mp$H_2$MM identified to capture dynamic interconversion (stars) were indeed found well above the static line, implying dynamics. f) apo correction of extracted rates. The apo contribution is corrected for by finding the minimum difference between measured rates (panel d) and the numerical mixing of apo values with a range of opening and closing rates. The white '+' marks the recovered dwell times, i.e. the position of the lowest residuals.

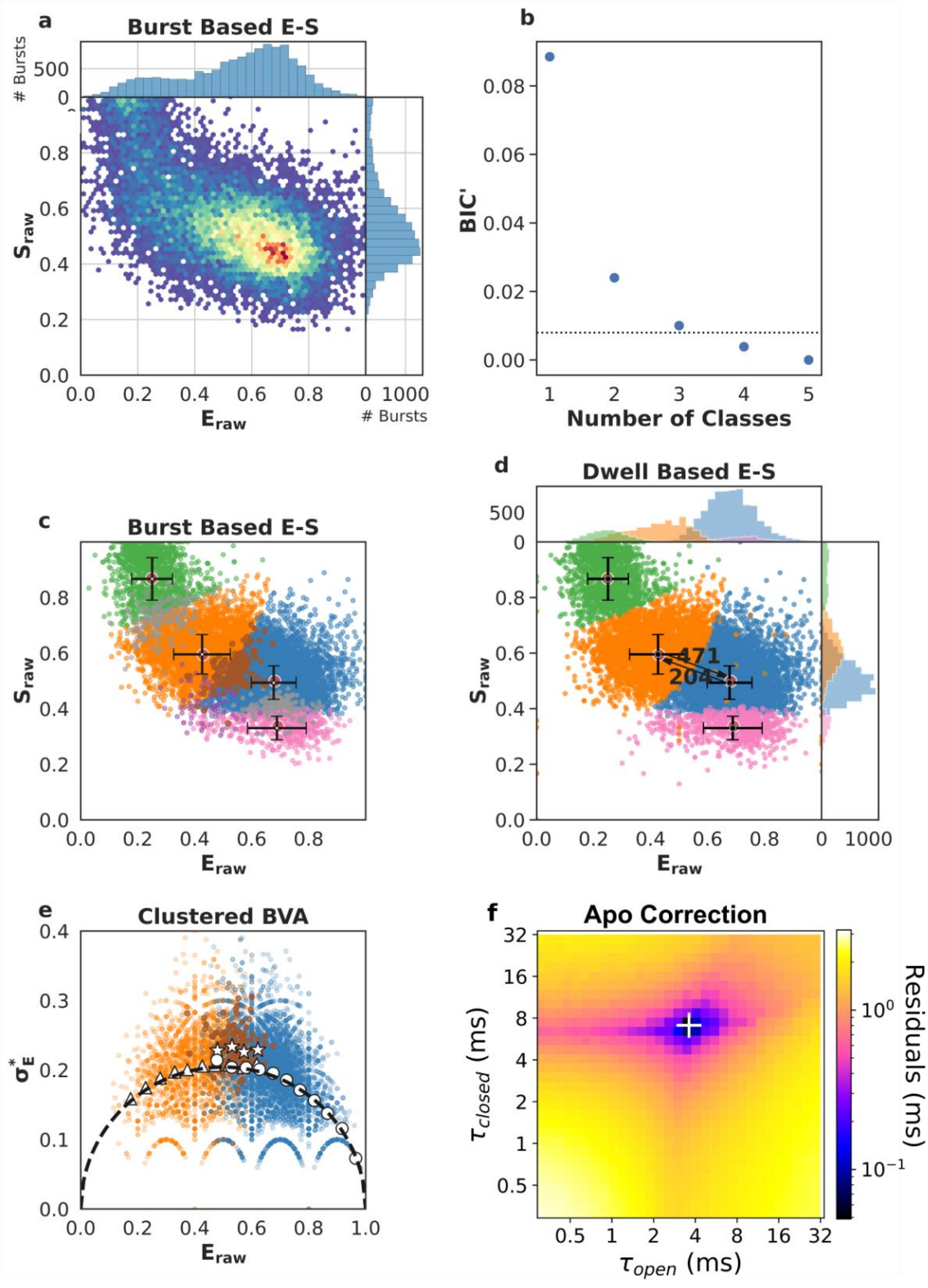
a
Burst Based E-S
### Bursts
500
0
0.8
0.6
0.4
0.2
0.0
S_raw
0.0 0.2 0.4 0.6 0.8
E_raw
0 1000
### Bursts
b
0.08
0.06
0.04
0.02
0.00
BIC'
1 2 3 4 5
Number of Classes
c
Burst Based E-S
d
Dwell Based E-S
471
204
e
Clustered BVA
σ*_E
1.0
f
Apo Correction
τ_closed (ms)
τ_open (ms)
Residuals (ms)
32 16 8 4 2 1 0.5

**Supplementary Fig. 15** - Single-molecule data for one of five repeats of SecYEG-PrlA4 + proSpy + ATP (SecYEG-PrlA4:A:ATP+proSpy) (first column of data in Supplementary Table 6 and 7). a) Burst data plotting FRET efficiency ($E_{raw}$) against stoichiometry ($S_{raw}$). b) Modified Bayes Information Criterion (BIC') of different state-models indicated that four classes best describe the data (most likely state model BIC'<0.005). c) Using a burst based $E_{raw}$ $S_{raw}$ plot we interpreted the classes as the open state (orange, low $E_{raw}$), closed state (blue, high $E_{raw}$), dark acceptor (green, high $S_{raw}$), dark donor (pink, low $S_{raw}$), in this step, we also identified bursts representing transitions between conformational states (brown), transitions and photophysics (purple) and pure photophysics (grey). Crosses identify the average $E_{raw}$ $S_{raw}$ position for each state and the standard deviations. d) Dwell based $E_{raw}$ $S_{raw}$ plot showing the rate of interconversion ($s^{-1}$) between the open and closed states. e) Burst Variance Analysis (BVA) provided a qualitative confirmation of dynamic exchange between states and transition detection within bursts. Binned values representing bursts assigned to either the open (triangles) or the closed (circles) state, respectively, coincide with the static line (black), exhibiting no exchange on a millisecond time scale. On the other hand, bursts which mp$H_2$MM identified to capture dynamic interconversion (stars) were indeed found well above the static line, implying dynamics. f) apo correction of extracted rates. The apo contribution is corrected for by finding the minimum difference between measured rates (panel d) and the numerical mixing of apo values with a range of opening and closing rates. The white '+' marks the recovered dwell times, i.e. the position of the lowest residuals.

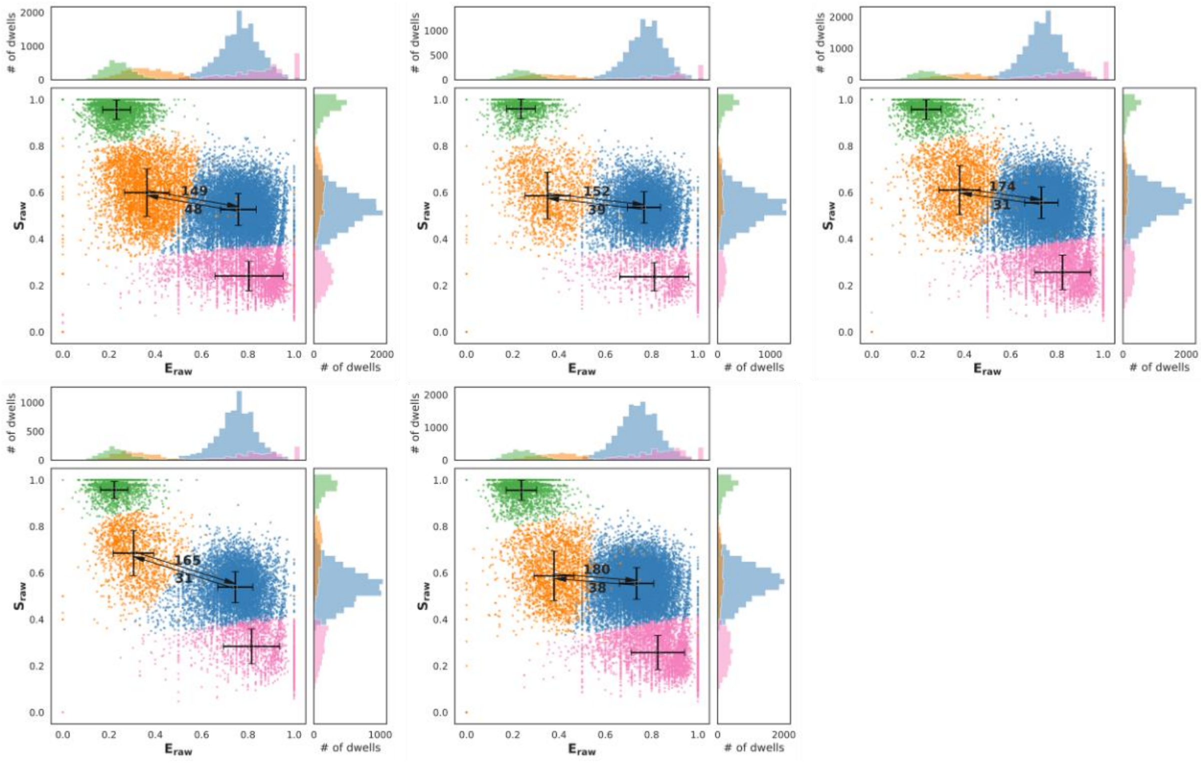


**Supplementary Fig. 16** - Five technical repeats show the variability of SecYEG channel opening and closure rates ($s^{-1}$) for wild-type SecYEG apo. The data for the SecYEG complex are obtained from a single protein purification, but proteoliposomes were prepared fresh before each measurement.

#### Supplementary Tables

| Condition | FRET State | Dwell Time (ms) | | | | | Mean Dwell Time (ms) ± 90%CI |
|---|---|---|---|---|---|---|---|
| SecYEG (apo) | Open | 6.7 | 6.6 | 5.7 | 6.1 | 5.6 | 6.1 ± 0.5 |
| | Closed | 20.8 | 25.6 | 32.3 | 32.3 | 26.3 | 27.5 ± 4.6 |
| SecYEG+SS (SS) | Open | 2.1 | 6.5 | 3.9 | 10.2 | 4.4 | 5.4 ± 2.9 |
| | Closed | 2.2 | 1.1 | 1.6 | 1 | 0.6 | 1.3 ± 0.6 |
| SecYEG:A (SecA) | Open | 2.5 | 1.9 | 4.2 | 1.3 | 2.6 | 2.5 ± 1.0 |
| | Closed | 40 | 32.1 | 19.5 | 34.5 | 15.3 | 28.3 ± 10 |
| SecYEG:A:ATPγS (ATPγS) | Open | 0.9 | 0.3 | 0.4 | 0.3 | 0.2 | 0.4 ± 0.3 |
| | Closed | 0.7 | 1.0 | 0.9 | 1.2 | 1.5 | 1.1 ± 0.3 |
| SecYEG:A:ADP·AlFx (ADP·AlFx) | Open | 1 | 3.7 | 4.5 | 4.3 | 4.0 | 3.5 ± 1.3 |
| | Closed | 1.6 | 2.0 | 1.9 | 2.1 | 1.9 | 1.9 ± 0.2 |
| SecYEG:A:ADP (ADP) | Open | 0.2 | 0.6 | 1.0 | 0.4 | 0.5 | 0.5 ± 0.3 |
| | Closed | 29.0 | 20.0 | 10.0 | 8.3 | 5.0 | 14.5 ± 9.4 |
| SecYEG:A:ATP (ATP) | Open | 2.0 | 2.3 | 1.3 | 3.9 | 4.5 | 2.8 ± 1.3 |
| | Closed | 2.5 | 2.2 | 3.0 | 2.0 | 2.4 | 2.4 ± 0.4 |
| SecYEG:A:ATP+proSpy (ATP+proSpy) | Open | 1.2 | 1.1 | 1.2 | 2.5 | 1.1 | 1.4 ± 0.6 |
| | Closed | 1.2 | 1.1 | 1.2 | 2.5 | 1.1 | 1.4 ± 0.5 |

**Supplementary Table 1** – SecYEG wild-type corrected dwell times in each FRET state (open = low FRET, closed = high FRET). Each experimental condition was repeated five times with fresh proteoliposomes.  Error is given as the 90% confidence interval (± 90%CI).

| Condition | Percent Open | | | | | Mean Percent Open ± 90%CI |
|---|---|---|---|---|---|---|
| SecYEG (apo) | 24 | 20 | 15 | 16 | 17 | 19 ± 4 |
| SecYEG+SS (SS) | 49 | 86 | 71 | 91 | 88 | 77 ± 17 |
| SecYEG:A (SecA) | 6 | 6 | 18 | 4 | 15 | 9 ± 6 |
| SecYEG:A:ATPγS (ATPγS) | 56 | 23 | 31 | 20 | 12 | 29 ± 16 |
| SecYEG:A:ADP·AlFx (ADP·AlFx) | 40 | 65 | 70 | 67 | 68 | 62 ± 12 |
| SecYEG:A:ADP (ADP) | 1 | 3 | 9 | 5 | 9 | 5 ± 4 |
| SecYEG:A:ATP (ATP) | 44 | 51 | 30 | 66 | 65 | 51 ± 14 |
| SecYEG:A:ATP+proSpy (ATP+proSpy) | 47 | 46 | 44 | 64 | 37 | 48 ± 9 |

**Supplementary Table 2** – SecYEG wild-type percent open ($\tau_{open}/(\tau_{open}+\tau_{closed})$x100, data from Supplementary Table 1). Each experimental condition was repeated five times with fresh proteoliposomes. Error is given as the 90% confidence interval (± 90%CI).

| SecYEG | proSpy | SecA ATPase Rate ($k_{cat}$) ($s^{-1}$) | | | | Mean SecA ATPase Rate ($k_{cat}$) ($s^{-1}$) ± 95%CI |
|---|---|---|---|---|---|---|
| Wild-type | + | 6.12 | 6.46 | 7.22 | 6.27 | 6.52 ± 0.42 |
| | - | 0.20 | 0.12 | 0.12 | 0.15 | 0.15 ± 0.02 |
| PrlA4 | + | 6.46 | 7.61 | 7.26 | 6.86 | 7.05 ± 0.43 |
| | - | 0.19 | 0.20 | 0.15 | 0.17 | 0.18 ± 0.01 |

**Supplementary Table 3** – The SecA ATPase rate ± proSpy in the presence of SecYEG proteoliposomes (wild-type and prlA4) and ATP. Each condition was performed with 4 technical repeats. Error is given as the 95% confidence interval (± 95%CI).

| SecYEG | Pep86 Position | Amino Acids | Lag Time (s) | | |
|---|---|---|---|---|---|
| Wild-type | 1 | 146 | 41.2 | 41.4 | 42.3 |
| | 2 | 292 | 91.2 | 90.7 | 93.9 |
| | 3 | 438 | 131.8 | 131.6 | 137.4 |
| | 4 | 584 | 196.3 | 189.0 | 209.7 |
| PrlA4 | 1 | 146 | 10.4 | 11.3 | 15.3 |
| | 2 | 292 | 25.0 | 26.8 | 31.7 |
| | 3 | 438 | 36.8 | 37.0 | 40.4 |
| | 4 | 584 | 53.6 | 55.4 | 68.1 |

**Supplementary Table 4 –** proSpy$_{4x}$ transport assay lag times using the NanoLuc split-luciferase assay for wild-type and PrlA4 SecYEG (for further details on the NanoLuc split-luciferase assay see Supplementary Methods). Each condition was performed with 4 technical repeats.

| SecYEG | Slope (± SE) | Intercept (± SE) | Coefficient of determination ($R^2$) | Transport Rate ($aa.s^{-1}$) (± SE) |
|---|---|---|---|---|
| Wild-type | 0.35 ± 0.01 | -11.59 ± 5.28 | 0.99 | 2.85 ± 0.03 |
| PrlA4 | 0.10 ± 0.01 | -3.30 ± 3.41 | 0.94 | 9.70 ± 0.02 |

**Supplementary Table 5** - Fitting statistics for Nanoluc assay lag times of $proSpy_{4x}$ (linear least-squares regression fits in Fig. 5a, data in Supplementary Table 4) and the derived transport rates (1/Slope) for wild-type and PrlA4 SecYEG. Error is given as standard error (± SE).

| Condition | FRET State | Dwell Time (ms) | | | | | Mean Dwell Time (ms) ± 90%CI |
|---|---|---|---|---|---|---|---|
| SecYEG-PrlA4 (apo) | Open | 1.5 | 1.3 | 2.0 | 1.2 | 1.6 | 1.5 ± 0.3 |
| | Closed | 4.2 | 3.6 | 7.5 | 2.4 | 4.9 | 4.5 ± 1.8 |
| SecYEG-PrlA4+SS (SS) | Open | 1.3 | 2.5 | 2.9 | 2.8 | 2.4 | 2.4 ± 0.6 |
| | Closed | 1.2 | 1.7 | 1.5 | 1.7 | 1.7 | 1.6 ± 0.2 |
| SecYEG-PrlA4:A (SecA) | Open | 2.3 | 3.0 | 2.8 | 3.3 | 2.4 | 2.8 ± 0.4 |
| | Closed | 1.5 | 1.4 | 1.7 | 1.5 | 1.4 | 1.5 ± 0.1 |
| SecYEG-PrlA4:A:ATP+proSpy (ATP+proSpy) | Open | 3.9 | 6.1 | 5.9 | 2.4 | 4.3 | 4.5 ± 1.5 |
| | Closed | 7.9 | 5.1 | 14.2 | 6.6 | 7.7 | 8.3 ± 3.3 |

**Supplementary Table 6 –** SecYEG-PrlA4 corrected dwell times in each FRET state (Open = low FRET, Closed = high FRET). Each experimental condition was repeated five times with fresh proteoliposomes. Error is given as the 90% confidence interval (± 90%CI).

| Condition | Percent Open | | | | | Percent Open ± 90%CI |
|---|---|---|---|---|---|---|
| SecYEG-PrlA4 (apo) | 26 | 26 | 21 | 34 | 25 | 26 ± 4 |
| SecYEG-PrlA4+SS (SS) | 52 | 60 | 66 | 62 | 59 | 60 ± 6 |
| SecYEG-PrlA4:A (SecA) | 61 | 68 | 62 | 69 | 63 | 65 ± 4 |
| SecYEG-PrlA4:A:ATP+proSpy (ATP+proSpy) | 33 | 54 | 29 | 27 | 36 | 36 ± 10 |

**Supplementary Table 7** – SecYEG-PrlA4 percent open ($\tau_{open}/(\tau_{open}+\tau_{closed})$x100, data from Supplementary Table 6). Each experimental condition was repeated five times with freshly prepared proteoliposomes. Error is given as the 90% confidence interval (± 90%CI).
